# Genome maintenance functions of *Trypanosoma brucei* DNA Polymerase N include telomere association and a role in antigenic variation

**DOI:** 10.1101/682948

**Authors:** Andrea Zurita Leal, Marie Schwebs, Emma Briggs, Helena Reis, Leandro Lemgruber, Katarina Luko, Falk Butter, Richard McCulloch, Christian J. Janzen

## Abstract

Maintenance of genome integrity is critical to guarantee transfer of an intact genome from parent to offspring during cell division. DNA polymerases (Pols) provide roles in both replication of the genome and the repair of a wide range of lesions. Amongst replicative DNA Pols, translesion DNA Pols play a particular role: replication to bypass DNA damage, often at the cost of mutation. All cells express a range of translesion Pols, but little work has examined their function in parasites, including whether the enzymes might contribute to host-parasite interactions. Here, we describe a dual function of translesion PolN in African trypanosomes. Previously we demonstrated that PolN is associated with telomeric sequences and now we show that RNAi-mediated depletion of PolN results in slowed growth, altered DNA content, changes in cell morphology, and increased sensitivity to DNA damaging agents. Depletion of PolN leads to chromosome segregation defects and accumulation of DNA damage. We also show that PolN displays discrete localisation at the nuclear periphery in the absence of exogenous DNA damage. In addition, we demonstrate that PolN depletion leads to deregulation of telomeric variant surface glycoprotein genes, linking the function of this translesion DNA polymerase to host immune evasion by antigenic variation.

## Introduction

Accurate duplication of the genome is a critical component of the cell cycle of all organisms. Two pathways contribute to accurate genome duplication: copying of the genome, and repair of DNA damage. Eukaryotic cells encode a wide range of DNA polymerases (Pols) that are required for DNA synthesis, allowing genome duplication, and for repair of DNA damage (review in [1]). Eukaryotic DNA Pols are divided into four different families (A, B, X and Y) based on sequence and structural homologies. Nuclear DNA Pols that direct the accurate copying of the genome belong to the B family, while mitochondrial genome replication is catalysed by an A family DNA Pol [2]. DNA Pols that act in DNA repair span all families, as do so-called translesion DNA Pols, which straddle DNA repair and replication activities because their activity is required whenever replicative DNA Pols encounter lesions in the template strand that must be bypassed to allow genome duplication [3];[4];[5]. In general, DNA replication is a high fidelity process with an extremely low error rate [6]. This is due to a combination of the ability of replicative DNA Pols to efficiently select the correct nucleotide to incorporate into the newly synthesized DNA strand and proofreading activity of the Pols, which permits the excision of occasionally incorrectly inserted nucleotides. Additionally, postreplicative repair mechanisms further reduce overall error rates by removing mispaired or damaged bases [7]. Although the wide range of DNA repair mechanisms available to all cells can efficiently detect and remove a myriad of lesions from the DNA template, some forms of lesions persist and risk the survival of the cell because an unrepaired lesion can lead to replication fork stalling and, potentially, death [8];[9]. Translesion synthesis (TLS) circumvents this problem [7], using TLS Pols to insert nucleotides in the new DNA strand and thereby bypassing a lesion in the template DNA strand. Recruitment of TLS Pols to damaged DNA is mediated by the proliferating cell nuclear antigen, PCNA [10]. The homotrimeric PCNA complex encircles DNA and interacts with replicative DNA Pols, increasing their processivity [11]. PCNA also interacts with TLS Pols through a PIP box motif [12]. Indeed, it has been suggested that at least some TLS Pols form a multi-protein complex at stalled replication forks [13]. Replication fork stalling also causes a prolongation of singlestranded DNA, which is recognized by the replication protein A (RPA) heterotrimer. RPA binding triggers mono-ubiquitination of PCNA by the RAD18/RAD6 complex [14], which facilitates the exchange of replicative polymerases with TLS polymerases and, thus, the bypass of a DNA lesion during replication.

Very little is known about TLS activity in *Trypanosoma brucei*, the causative agent of African Sleeping Sickness in humans and the devastating animal disease *Nagana* in sub-Saharan Africa. The only functional study to date described two primase-polymerase-like proteins called PPL1 and PPL2 [15]. TLS activity of both polymerases was confirmed by their ability to insert nucleotides opposite thymine dimers in DNA templates *in vitro*. Moreover, depletion of PPL2 causes a severe cell cycle defect after most nuclear DNA synthesis is complete and activation of the DNA damage response [15]. What endogenous feature(s) of the nuclear *T. brucei* genome PPL2 acts upon is unknown, though the TLS Pol was very recently shown to be a component of telomere-binding protein complexes in trypanosomes [16]. Telomere-associated proteins are of particular interest in *T. brucei*. Besides their function in telomere maintenance, they are involved in one of the most important strategies used by pathogens to evade the vertebrate host immune response: antigenic variation (reviewed in [17]). In *T. brucei*, antigenic variation is mediated by a periodic exchange of a protective coat composed of variant surface glycoprotein (VSG) [18]. There are more than 2000 VSG genes or pseudogenes in the *T. brucei* genome [19];[20] but at any given time only a single VSG gene is expressed from one of approximately 15 specialized loci, the so-called bloodstream VSG expression sites (BES) [21]. BES are always located adjacent to telomeres and switching of the expressed VSG can occur by events that change which of the BES is singularly transcribed, or by recombination reactions that replace the BES VSG with a silent gene [22]. Several telomere-associated proteins are known to be involved in the transcriptional control mechanisms that ensure only a single BES is transcribed, and to affect VSG recombination [17]. For example, repressor activator protein 1 (RAP1) appears to be necessary for monoallelic expression of VSG genes because depletion of RAP1 partially de-represses all silent BES [23]. Furthermore, it was shown that the telomere duplex DNA-binding factor TRF suppresses homologous recombination of VSG sequences into the active BES [24]. The TRF interacting factor 2 (TIF2) also appears to be a negative regulator of VSG switching in collaboration with TRF, but by an additional mechanism that is independent from TRF association [25];[26]. Beyond such telomere-focused observations, recent work has shown that the actively transcribed BES is replicated early in S-phase, whereas all silent BES are replicated later [27]. What factors dictate that only the transcribed BES is early replicating, and how the BES replication process relates to the rest of the genome, is unknown [28];[29];[30]. Furthermore, accumulating evidence indicates that the active BES contains RNA-DNA hybrids [31];[32], which can lead to DNA damage, and all BES are prone to lesions [33], but whether these observations relate to specific aspects of BES replication is unknown.

Since the exact mechanisms of BES monoallelic transcriptional control and potential targeting of recombination events in the BES remain elusive, we were intrigued to find that, in addition to PPL2, a further putative TLS Pol, termed PolN, is a telomere-associated protein, since it could be purified with a telomeric repeat-containing oligonucleotide and by co-immunoprecipitation with TRF [16]. Here, we show that PolN displays discrete localisation to the nuclear periphery of *T. brucei* mammal-infective cells and that RNAi-mediated depletion of PolN results in slowed growth without a specific cell cycle arrest, accumulation of DNA damage and chromosome segregation defects. We also document substantial deregulation of telomeric VSG genes after PolN depletion, which suggests at least one TLS DNA Pol can contribute to antigenic variation.

## Material and Methods

### Sequence analysis

*T. brucei* gene and protein sequences were retrieved from TriTrypDB version 33 (http://tritrypdb.org/tritrypdb/). Multiple alignment sequence analysis was conducted using ClustalW. TbPolN protein domain analysis was performed using InterPro 64.0 (https://www.ebi.ac.uk/interpro/) and Pfam 31.0 (http://pfam.xfam.org/).

### Maintenance of cell cultures

Lister 427 (WT) bloodstream form (BSF) cells were grown in HMI-9 (Gibco®) (Hirumi and Hirumi, 1989), supplemented with 20% (v/v) heat inactivated fetal bovine serum (FBS; Sigma Aldrich) and 1% penicillin-streptomycin solution (stock at 10,000 U/ml) (Gibco®). In the case of the 2T1 cell line, the cells were grown in HMI-11 thymidine-free media, consisting of Iscove’s Modified Dulbecco’s Medium (IMDM) (Gibco®), 10% (v/v) of FBS (Gibco®,tetracycline free), 1% of penicillin-streptomycin solution (10,000 U/ml)(Gibco®). For the RNAi cell lines used in EdU replication assays, the cells were grown in HMI-11 thymidine free media, consisting of Iscove’s Modified Dulbecco’s Medium (IMDM) (Gibco®), 10% (v/v) of FBS (Gibco®, tetracycline free), 1% of penicillin-streptomycin solution (10,000 U/ml; Gibco®), 4% (v/v) of HMI-9 mix (0.05 mM of bathocuproine disulphonic acid, 1 mM of sodium pyruvate, and 1.5 mM of L-cysteine, 1 mM of hypoxanthine and 0.0014% of 2-mercaptoethanol (Sigma Aldrich)). For Lister 427 (WT) no drugs were added to the media. The selective drugs used for 2T1 cells were puromycin (0.2 µg/ml) and phleomycin (2.5 µg/ml), for RNAi cell lines phleomycin (2.5 µg/ml) and hygromycin (5µg/ml) and for N50 [34] neomycin (2.5 µg/ml). The tagged cells were grown in medium with 10 µg/ml blasticidin.

### Western blot

Western botting was performed according standard protocols, using approximately 2×10^6^ parasites. TbPolN-12myc was detected using mouse anti-myc clone 4A6 antibody at 1:7000. The detection of Ef1α (loading control), was performed with the mouse anti-Ef1α clone CBP-KK1 antibody at 1:25000. Both antibodies were used in combination with the goat anti-mouse IgG (H+L) horseradish peroxidase conjugate at 1:3000. While the damage accumulation marker γH2A was detected using rabbit anti-γH2A at 1:10000 in combination with goat anti-rabbit IgG (H+L) horseradish peroxidase conjugate at 1:3000. To analyse VSG expression, cell lysates of approximately 2×10^6^ cells were separated by SDS-PAGE and transferred to a PVDF membrane. VSGs were detected using polyclonal antibodies from rabbits immunized with VSG2, VSG3, VSG6 or VSG13. Signal intensity was normalized using a monoclonal antibody L13D6 specific for a paraflagellar rod protein (Gift from Keith Gull, University of Oxford). IRDye680- or IRDye800-conjugated secondary antibodies were used in combination with an Odyssey infrared scanner (LI-COR).

### TbPolN endogenous epitope tagging

TbPolN gene was modified *in situ* in *T. brucei* BSF cells, strain Lister 427, in order to express the protein as a C-terminal fusion of the myc epitope. To accomplish this, the protein was C-terminal tagged with 12 tandem repeats of the myc epitope using the pNAT^12myc^ BSD vector [35]. The 3’ end of TbPolN ORF, with the exclusion of the stop-codon, was PCR-amplified using the following primers Fw-gcatgagctcacgagttgctcattaagcac and Rv-gcattctagaaggaacatcaagtttctcga. The forward primer contained a SacI restriction site and the reverse primer contained a XbaI restriction site, facilitating the cloning of the fragment into the vector. The resulting vector was linearized prior to transfection using PstI. The transformants were selected using 10 µg/ml blasticidin.

### TbPolN cellular localization

TbPolN cellular localization was tested before and after genotoxic stress. Two cultures containing 2×10^5^ cells were grown in the presence of 10 µg/ml basticidin, one culture was treated with 0.003% MMS. After 18 hrs, 2×10^6^ parasites were harvested by centrifugation (405x g) and washed with PBS. Next, 25 µl of the cell suspension was loaded to 12 multi-well glass slide, pre-treated with Poly-L-lysine and allowed to settle for 5 min. The cells were fixed with 4% paraformaldehyde for 5 min and permeabilised with 0.2 % Triton X-100 (Promega) in PBS for 10 min. Next, 100 mM glycine was added for 20 min and washed twice with PBS for 5 min. Afterwards, the cells were incubated with 1% BSA/0.2% Tween-20 in PBS for 1 hr. The mouse anti-myc 4A6 Alexa Fluor® 488 conjugated antiserum diluted 1:500 in 1% BSA/0.2% Tween in PBS was applied for 1hr, followed by two washes with 3% BSA in PBS. After, the Fluoromount G containing DAPI mounting medium was added; the slide was covered with a coverslip and sealed with regular nail varnish. The images were captured in the Deltavision RT deconvolution fluorescence microscope, the 1.4/63 x lens was used. Z-stacks of 6 µm (30 sections, 0.2 µm thickness each) were acquired using SoftWoRx Suite 2.0 (Applied Precision, GE). High resolution images were captured on an Elyra PS.1 super resolution microscope (Zeiss) using the 1.4/63 x lens. The images were acquired using the ZEN Black Edition Imaging software.

### Construction of TbPolN RNAi cell line

A fragment of the TbPolN gene was PCR amplified from *T. brucei* TREU 927, using primers PolNC1, PolNC4, PolNC9 (sequences upon request), incorporating the attB1 and attB2 sites. The fragment was cloned into the pPGL2084 vector [36] in a BP recombinase reaction, using the Gateway® BP clonase® II Enzyme mix Kit (Life Technologies), following the manufacturer’s instructions. The final vector was linearized with AscI and transfected into the genetically modified Lister 427 strain, 2T1. The transformants were selected with 5 µg/ml hygromycin, in the presence of 2.5 µg/ml phleomycin.

### Survival assay

TbPolN RNAi cell line was cultured in tetracycline (Tet) free medium containing phleomycin and hygromycin. Parasites were dilute into two cultures of 1×10^4^ cells/ml, one containing 1 µg/ml tetracyclin (Tet+) to induce the RNAi-mediated depletion of the gene. The population density was assessed every 24 for a period of 3 days, using the Neubauer haemocytometer. The cells survival was also tested after the exposure to genotoxic stress. The cells were prepared as described above. Both flasks were mixed and 1ml of each culture aliquoted to a 24 well plate. The appropriate concentration (0%, 0.001%, 0.002%, 0.003%) of MMS was added to each well of Tet- and Tet+ cells, assessing the population density every 24 hrs. In the case of UV exposure the Tet- and Tet+ cells were UV irradiated (0 J/m^2^, 500 J/m^2^, 750 J/m^2^, 1000 J/m^2^ and 1500 J/m^2^) 24 hrs post induction using a Stratagene Stratalinker UV Crosslinker 2400.

### Damage accumulation assay

Two cultures containing 2×10^4^ *Tb*PolN RNAi cells were grown in the absence and presence of 1 µg/ml of tetracycline. After 24 hrs, 48 hrs and 72 hrs, 2×10^6^ parasites were harvested by centrifugation (405x g) and washed with 1x PBS. The cells were prepared for western blot and immunofluorescence as described above. For both approaches anti-γH2A antibody, which recognises Thr130 phosphorylated histone H2A, was diluted to a concentration of 1:1000 and detected with α-rabbit HRP conjugate (1:3000) for western blot and Alexa Fluor® 594 α-rabbit (1:1000) for immunofluorescence analysis.

### EdU incorporation assay

TbPolN RNAi cells were incubated with 150 µM of 5-ethynil-2’-deoxyuridine (EdU; Life Technologies) for 4 hrs at 37 °C with 5% CO_2_. The cells were then harvested by centrifugation at 1000x g for 5 min and washed with 1x PBS. The pellet was re-suspended in 20 µl of 1x PBS and the cell suspension was pipetted onto a 12 well slide, pre-treated with poly-L-lysine, and left to settle for 5 min. The supernatant was removed and 25 µl of 4% paraformaldehyde was added and left for 4 min, followed by a wash with 3% BSA in 1x PBS. The cells were permeabilised by adding 20 µl of 0.2% Triton X-100 (Promega) in 1x PBS for 10 min at room temperature. The wells were washed twice with 3% BSA in 1x PBS. The supernatant was removed and 25 µl of the Click-iT® (Life Technologies) reaction, containing 21.25 µl of 1x Reaction Buffer, 1 µl of CuSO_4_, 0.25 µl Alexa Fluor® 555 Azide and 2.5 µl of 1x Reaction Buffer, was added for 1 hr at room temperature. The solution was removed and the cells were washed 6 times with 3% BSA in 1x PBS. The Fluoromount G containing DAPI mounting medium was added and the slide was covered with a coverslip and sealed. In the case of a double labelling detection, EdU signal and phosphorylated γH2A signal, the sample was incubated first with the Click-iT® reaction, followed by 1 hr incubation with 1% BSA/0.2% Tween-20 in 1x PBS. Afterwards, anti-γH2A antibody diluted 1:1000 was added for 1 hr and detected with Alexa Fluor® 594 α-rabbit diluted 1:1000. The images were captured using a Zeiss Axioskop 2 fluorescent microscope (63x/ 1.4 oil objective). The images were acquired using the ZEN software package (Zeiss; http://www.zeiss.com/corporate/en_de/home.html).

### Measurement of fluorescence intensity

For the intensity analysis the software Fiji (ImageJ) was used to convert images to grayscale and to subtract the background. A circle of 2.1 x 2.1 pixel region of interest (ROI) was drawn around the cell, obtaining the density (area and the mean gray value) and mean fluorescence value of each cell. For each image a region with no fluorescence was selected, which corresponds to the background control. The fluorescence intensity was obtained by calculating the corrected total cell fluorescence (CTCF) using the following formula:

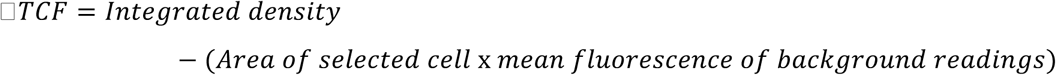

### Fluorescence *in situ* hybridisation (FISH)

Approximately 5×10^6^ cells were harvested by centrifugation at 405x g for 10 min and washed in 1x PBS at 660x g for 3 min. The supernatant was removed and the pellet re-suspended in 4% formaldehyde in 1x PBS for 20 min at room temperature. The cells were washed and re-suspended in 65 µl of 1x PBS and spread onto a pre-treated poly-L-lysine (Sigma Aldrich) slide and left to settle for 20 min. The supernatant was removed and the cells permeabilised with 0.1 % TritonX-100 in 1x PBS for 3 min. The cells were washed and dehydrated with pre-chilled ethanol in ascending concentration (70-90-100%) for 5 min each concentration. The slide was left to dry. In the meantime, 10 µl of the telomere probe (TTAGGG; Thermo Fisher) was added into 60 µl of hybridization solution (50 % Formamide, 10% Dextran, 2x SSPE buffer [1x SSPE: 0.18 M NaCl, 10 mM NaH_2_P0_4_, 1 mM EDTA, pH 7.7: use buffer at pH 7.9]) and heated at 85 °C in a water bath for 7 min. Once the slide was completely dry the hybridization solution containing the probe was added. The slide was covered with a plastic coverslip and incubated at 95 °C for 5 min (water bath), followed by 16 hrs incubation at 37 °C. Then, the coverslip was removed and the slide was washed for 30 min with 2x SSC (Thermo Fisher)/50% Formamide solution at 37 °C, 60 min with 0.2x SCC at 50 °C and a final wash with 4x SSC at room temperature for 10 min. The slide was finally mounted in Fluoromount G containing DAPI and sealed with nail varnish.

### Cell viability assay

Approximately 1×10^6^ cells were harvested and washed twice with 1 ml TDB (1500xg, 4°C, 10 min). The supernatant was discarded and the pellet was taken up in 400 µl TDB. Propidium iodide was added in a final concentration of 2,5 µg/ml to the resuspended pellet and incubated for 10 min on ice. Analysis of the dyed cells was performed with a BD FacsCalibur. For the analysis of dead cells, WT cells without puromycin resistance were incubated overnight with a final concentration of 1 µg/ml puromycin.

### Cell cycle analysis with propidium iodide

2×10^7^ cells harvested and washed with 1 ml PBS (1500xg, 4°C, 10 min). The cell pellet was resuspended in 2 ml ice-cold PBS/2 mM EDTA and fixed by the dropwise addition of 2,5 ml ice-cold EtOH. After 1 hr of incubation at 4°C the cells were washed with 1 ml PBS (1500xg, 4°C, 10 min) and the pellet resuspended in 1 ml PBS. 10 µg RNaseA and propidium iodide in a final concentration of 10 µg/ml were added to the suspension. After inverting of the sample three time, the cell suspension was incubated for 30 min at 37°C. Cells were stored at 4°C before analysis with a BD FACSCalibur.

### VSG-2 analysis by flow cytometry

All devices and solutions were precooled (4 °C). Approximately 1×10^6^ cells were harvested (1500 g, 4 °C, 10 min) and the cell pellet suspended in 100-500 µl HMI-9 medium. The VSG-2 CRD A antibody was added (1:500) and then the suspension was incubated rotating for 1 hr (6 rpm, 4 °C). The cells were sedimented and washed twice with 500 µl HMI-9 medium (1500 g, 4 °C, 10 min). The pellet was suspended in 500 µl HMI-9 medium and the secondary antibody, Alexa Flour® 488 anti-rat IgG, was added (1:2000). The suspension was incubated, with rotating for 10 min at 4 °C. The cells were sedimented, washed twice with 500 µl TDB (1500 g, 4 °C, 10 min) and resuspended in 400 µl TDB medium. The cells were stored at 4 °C until analysis using a BD FACSCalibur. Prior to this experiment, the procedure was conducted without addition of the primary and secondary antibodies. After resuspension of the pellet in 400 µl of TDB, propidium iodide was added to a final concentration of 2.5 µg/ml and incubated for 10 min on ice. At the BD FACSCalibur the gate was set such that only living cells were considered for the analysis.

### Isolation of soluble VSG for mass spectrometry analysis

Approximately 4×10^7^ cells were precooled for 10 min on ice and then harvested with 1500 xg for 10 min at 4 °C. The cell pellet was resuspended in 45 µl 10 mM sodium phosphate buffer with additional protease inhibitors (leupeptin, PMSF and TLCK). The suspension was incubated for 5 min at 37 °C and then cooled down for 2 min on ice. The cells were sedimented (14.000 g, 4 °C, 5 min) and only the supernatant transferred to a new reaction tube. 15 µl of 4x NuPage® Buffer (Invitrogen) supplemented with 400 mM DTT was added to the suspension and incubated at 70 °C for 10 min. The cell suspension was stored at −20 °C until the shipping to Falk Butter (IMB Mainz) for mass spectrometry analysis. The experiment was conducted in quadruplicate. The samples were loaded into a 10% NuPage NOVEX gel and run in 1x MES (Thermo) at 180V for 10 min. The gel was fixated and stained with Coomassie (Sigma). The gel lanes were sliced, destained in 50% EtOH/25 mM ABC (ammonium bicarbonate) pH 8.0, reduced in 10 mM DTT (Sigma) for 30 min at 56 degC and subsequently alkylated with 55 mM iodoacetamide (Sigma) for 30 min at RT in the darkd. Proteins were digested with MS-grade trypsin (Sigma) overnight at 37 degC and tryptic peptides desalted and stored on StageTips. For MS analysis, peptides were separated on an in-house packed Reprosil C18-AQ 1.9 µm resin (Dr. Maisch GmbH) capillary (New Objective) with an optimized 75 min gradient from 2% to 40% ACN (with 0.1% formic acid) prepared with an Easy-nLC 1000 system (Thermo). The packed capillary was mounted in a column oven (Sonation) and sprayed peptides continuously into a Q Exactive Plus mass spectrometer (Thermo) operated with a top10 data-dependent acquisition mode. Spray voltage was set between 1.8 −2.4 kV. The MS raw files were processed with MaxQuant (version 1.5.2.8) using standard settings with additionally activated LFQ quantitation and match between runs option. For the search, a concatenated database of TriTrypDB-8.1_TbruceiLister427_AnnotatedProteins.fasta (8,833 entries), TriTrypDB-8.1_TbruceiTREU927_AnnotatedProteins.fasta (11,567 entries) and the Lister427 VSG gene fasta (GAM Cross) translated in three reading frames vsgs_tb427_cds.out.fasta (86,764 entries) was used. For data analysis, all protein groups with less than 2 peptides (1 unique + 1 razor) and all protein groups not containing a VSG annotation as primary entry were removed. For the remaining Lister427 VSG proteins from the VSG gene fasta, the mean LFQ expression value of the quadruplicates was calculated and plotted as VSG abundance in R.

## Results

### T. brucei encodes separate DNA polymerase and helicase proteins related to the two domains found in eukaryotic PolQ (theta)

Eukaryotic cells, with the exception of fungi, express a range of factors related to DNA PolQ (theta), a protein that possesses both a C-terminal family A DNA Pol domain and an N-terminal super family 2 helicase domain (Fig.1, [37]). PolQ has been implicated in a wide range of cell functions, including lesion bypass [38];[39]; [40], DNA break repair by microhomology-mediated recombination (including during the repair of stalled DNA replication) [41];[42];[43];[44], and initiation of DNA replication [45]. In contrast, where organisms express related proteins possessing just the PolQ-like DNA Pol domain (PolN in mammals) or helicase domain (HelQ in mammals), their functions are less clearly understood [37]. PolN-like proteins have to date only been described in vertebrates, where TLS activities [46];[47];[48] and roles in inter-strand cross link and recombination repair [49];[50];[51] have been detailed, whereas HelQ-like proteins have been investigated in multicellular eukaryotes and archaea (Fig.1) [37];[52];[53];[54];[55]. In this evolutionary context, the PolQ-related protein complement found in *T. brucei* is highly unusual, since the parasite does not encode a PolQ protein with both DNA Pol and helicase domains, but encodes both PolN- and HelQ-like proteins (Fig.1). Thus, *T. brucei* and related trypanosomatids are the only organisms beyond mammals to possess genes encoding both truncated PolQ homologues, and the only organisms so far detailed that express only these proteins and not PolQ [56]. Whether this is because PolQ evolved in some eukaryotes as a gene fusion, or if *T. brucei* and relatives discarded PolQ is unclear. Nonetheless, the lack of *T. brucei* PolQ is perplexing, since the parasite displays robust microhomology-mediated end-joining activity [57];[58];[59].

**Figure 1.**
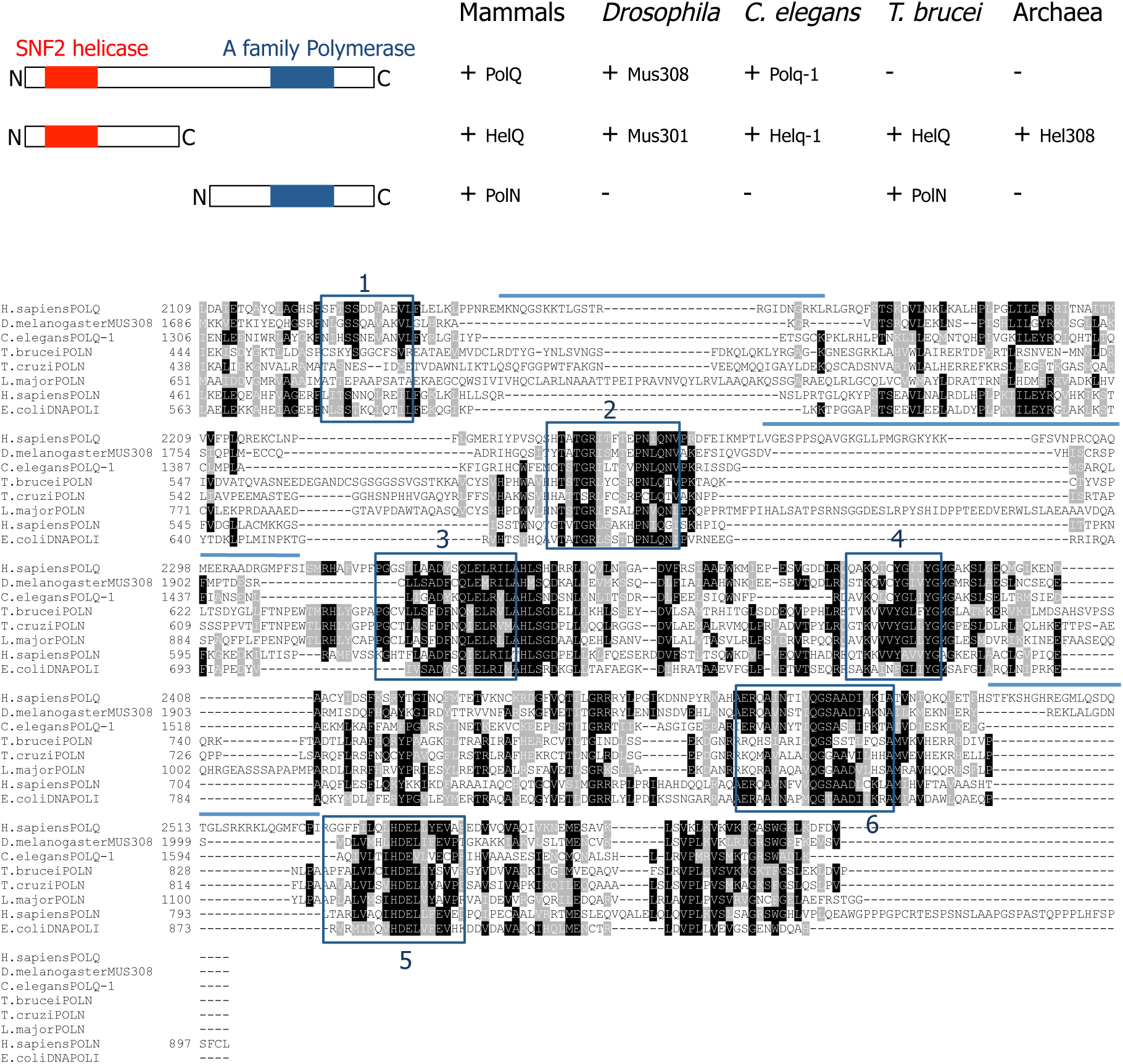
DNA Polymerase Q-related proteins in *Trypanosoma brucei*, selected multicellular eukaryotes and archaea. **A.** Schematic representation of the distribution of conserved SNF2 helicase (red) and family A (blue) polymerase (Pol) domains found in homologous proteins described in mammals, *Drosophila melanogaster*, *Caenorhabditis elegans*, *T. brucei* and archaea (protein names are provided). **B.** Sequence alignment of the Pol domain found in *H. sapiens* PolQ and PolN, *D. melanogaster,* Mus308, *C. elegans* Polq-1, *Escherichia coli* PolI and the predicted sequences of PolN from *T. brucei*, *T. cruzi* and *Leishmania major*. Boxes 1-6 show conserved family A Pol motifs, and lines illustrate inserts associated with lesion bypass activity of PolQ.

PolN-like genes are found syntenically in all kinetoplastids (Fig.1), and sequence alignments reveal the presence of six conserved Pol domains, consistent with Pol activity, as seen in purified protein from *Leishmania infantum* (there called PolQ, [60]). Intriguingly, sequence insertions between some of the Pol domains, which have been shown to underlie the lesion bypass activity of human PolQ [39], are much more marked in the *L. major* PolN than in the orthologues from *T. brucei* or *T. cruzi* (Fig.1). Whether this might mean alteration in TLS functions of the trypanosome proteins relative to the ability of *L. infantum* PolN to bypass 8oxoG, abasic sites and thymine glycol lesions *in vitro* is unclear [60].

### Super-resolution imaging reveals discrete subnuclear localisation of TbPolN

Localisation of *T. brucei* PolN (TbPolN) was assessed by immunofluorescence imaging using a cell line generated to express TbPolN as fusion to 12 C-terminal copies of the myc epitope (TbPolN-12myc; Fig.2). Western blotting revealed expression of a myc-tagged protein of the expected size (Fig.S1A). Intracellular localisation was expected to be nuclear [61] and examined with and without growth of the cells in the presence of the alkylating agent methyl methanesulfonate (MMS, 0.003%) for 18 hrs. In both conditions all anti-myc signal was found in the nucleus (Fig.S1B, C), and in the absence of MMS treatment confocal images suggested localisation of the protein in discrete subnuclear puncta (Fig.S1B). Structure illuminated super-resolution imaging revealed multiple puncta, with two relatively intense puncta present at the nuclear periphery in all cells (Fig.2). After growth in MMS, anti-myc signal was more dispersed throughout the nucleoplasm, perhaps suggesting redistribution of TbPolN-12myc in the presence of damage (Fig.2). What features of the nuclear genome or nucleus are predominantly localised in the two regions of the nuclear periphery is unclear. However, fluorescent imaging of both TbPolN-12myc and Thr130-phosphorylated histone H2A (gH2A), did not reveal extensive overlap of the two signals with or without MMS treatment (Fig.S2), suggesting TbPolN is not associated with the small numbers of relatively discrete lesions marked by gH2A in WT cells [62];[31], nor is it recruited to all the damage generated by MMS.

**Figure 2.**
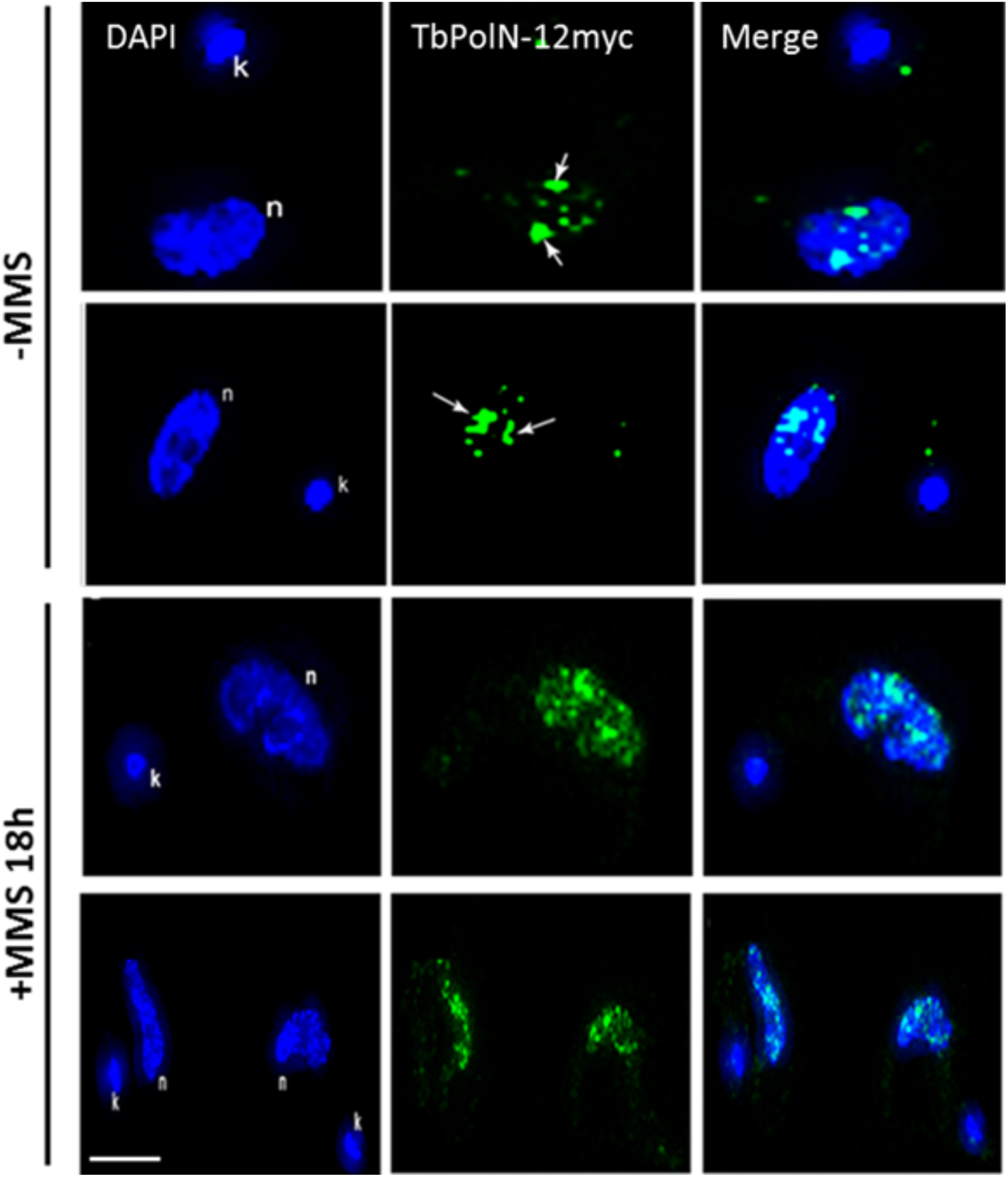
Subcellular localisation of TbPolN. Immunolocalisation of TbPolN-12myc in *T. brucei* BSF cells: left panels show DAPI staining of the nucleus (n) and kinetoplast (k), the middle panels show localisation of TbPolN-12myc using a conjugated Alexa Fluor® 488 anti-myc antibody, and the right panels are merged images of both signals. Images are shown of cells grown without damage (-MMS) or after growth for 18 hrs in the presence of 0.003% methyl methanesulphonate (+MMS 18 hrs). Arrows highlight two sites of significant TbPolN-12myc signal seen at the periphery of the nucleus in all untreated cells. Images were captured on a Zeiss Elyra Super Resolution Microscope. Scale bar represents 2 µm.

### PolN is important for growth of bloodstream form T. brucei

To learn more about the function of TbPolN in trypanosomes, we used tetracycline-inducible RNAi to deplete the protein in BSF cells (Fig. 3). Stagnation of growth was observed 24 h post induction of RNAi (Fig.3A). To ask if this growth defect could be explained by cell cycle arrest, we used flow cytometry to analyse the DNA content of propidium iodide-stained parasite populations at several time points after induction (Fig 3B). Numbers of 1N (G1 phase) cells were found to decrease in abundance with time, without equivalent accumulation of either 1N-2N (S phase) or 2N (G2/M) cells, suggesting there was no arrest at an observable cell cycle stage. Instead, loss of 1N cells was concomitant with increased numbers of cells with <1N and >2N DNA content, indicating that loss of TbPolN led to cells that had both lost and gained nuclear genome content. To ask if growth arrest after RNAi might also reflect cell death, we asked how many unpermeabilised cells could take up and be stained with propidium iodide (Fig. S3). 48 hrs after RNAi induction, when growth was strongly affected and substantial loss of G1 cells was detected, only around 10% of cells were classified as ‘dead’ by this approach. Hence, it seems unlikely that stagnation of cell growth after TbPolN loss is caused predominantly by cell death.

**Figure. 3.**
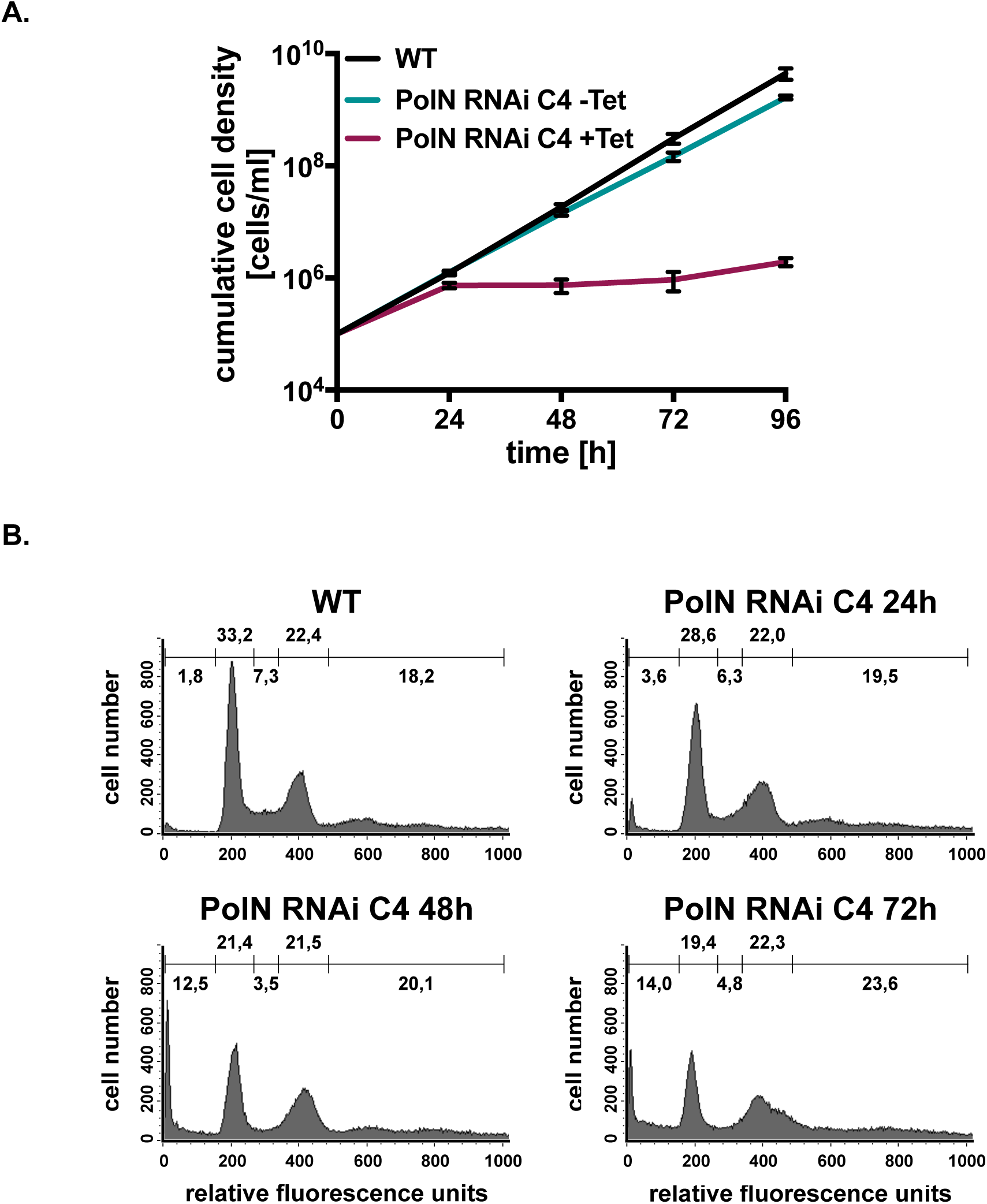
RNAi-depletion of TbPolN results in growth arrest. **A.** Cumulative cell density of the parental *T. brucei* RNAi strain 2T1 (WT), TbPolN RNAi cells grown without addition of tetracycline (Tet-), and TbPolN RNAi cells after induction of RNAi with tetracycline (Tet+). Error bars denote the standard deviation (*n* = 3). **B.** Cell cycle profiles of propidium iodide-stained parasites 24, 48 and 72 hrs after after RNAi-mediated depletion of PolN; parental 2T1 cells served as a control (WT). The values in the gates represent percentage of the whole population. Graphs display one representative analysis from three experimental repeats.

To ask if TbPolN contributes to DNA repair, growth was evaluated before and after RNAi in the presence of MMS and after exposure to UV light (Fig.S4). In each case, the extent of growth impairment caused by addition of the genotoxic agents was slightly more pronounced after RNAi than before. Thus, although TbPolN-12myc does not display extensive colocalisation with gH2A signal after MMS exposure (Fig.2), it may contribute to the response to alkylation damage and the distinct lesions caused by UV. Whether the range of damage TbPolN acts on overlaps with or is distinct from the orthologous protein in *L. infantum* is unclear, but LiPolN is also implicated in tackling a range of lesions [60], perhaps suggesting a wide range of activities for kinetoplastid PolNs.

### Analysis of aberrant cells 48-72 hrs post RNAi induction

To learn more about the effects of TbPolN depletion, cells from 24-72 hrs after RNAi induction were stained with 4’,6-diamidino-2-phenylindole (DAPI), which allows visualization of the trypanosome nucleus (N) and the kinetoplast (K) (Fig.4). Depending on the stage of the cell cycle the N-K content of an individual will vary, because replication and segregation of the nucleus and kinetoplast differ in their timing [63]. Cells displaying a 1N1K content are predominantly in G1, though some may have entered nuclear S (which can also be seen by an elongated K configuration and, perhaps, increased N staining). Cells with 1N2K configuration are predominantly in G2/M, and cells that have completed mitosis and are undergoing cytokinesis are 2N2K. Changes in the distribution of these configurations in mutants or after RNAi indicate altered cell cycle dynamics, and non-standard configurations have been detailed to arise after mutation or depletion of specific genes [64];[65]. In this experiment, RNAi was performed in cells in which TbPolN-12myc was expressed from its own locus, allowing us to evaluate the effectiveness of the knockdown. 24 hrs after depletion of TbPolN the DNA content distribution of the population and cell outline did not seem to be strongly affected, though a minor increase in the number of 1N2K cells, a small decrease in 2N2K cells and the appearance of cells with non-standard DNA configurations (‘other’) was apparent in light of the effects seen later. 48 hrs post RNAi induction a pronounced decrease in 1N1K cells and a further loss of 2N2K cells was clearly seen. In addition, though the numbers of 1N2K cells did not appear to increase further relative to 24 hrs, there was a very strong accumulation of cells with aberrant DNA content. At the same time, the cells had lost their typical trypomastigote shape. DNA content in the aberrant cells was hard to classify, but most cells had multiple kinetoplasts and abnormal, enlarged nuclei. These data suggest an initial, partial arrest in G2/M after loss of TbPolN, which is then overridden and the cells progress to mitosis but cannot complete it accurately, explaining the reduction in 2N2K cells, accumulation of abnormal cells, and the strong decrease of 1N1K (G1) cells. Overall, the DAPI analysis also appears consistent with the flow cytometry (Fig.3). Surprisingly, the DAPI time course also revealed that the pronounced cell cycle perturbations seen 48 hrs after RNAi induction were somewhat transient phenotypes, since at 72 hrs the numbers of 1N1K cells increased and the numbers of aberrant cells decreased relative to 48 hrs (Fig.4B); in addition, it was possible to observe greater numbers of cells with more regular body shape (Fig.4A). The basis for this change is unclear, since western blot analysis indicated that levels of TbPolN-12myc were reduced from 24 hrs after RNAi induction, and the protein was still undetectable at 72 hrs. Moreover, the apparent reversion was not associated with increased growth, as the RNAi cells still grew more slowly than uninduced controls at 72 hrs. Nonetheless, the shift from a severe to a less severe cell cycle perturbation from 48-72 hrs was mirrored in two other phenotypes we examined (see below).

**Figure 4.**
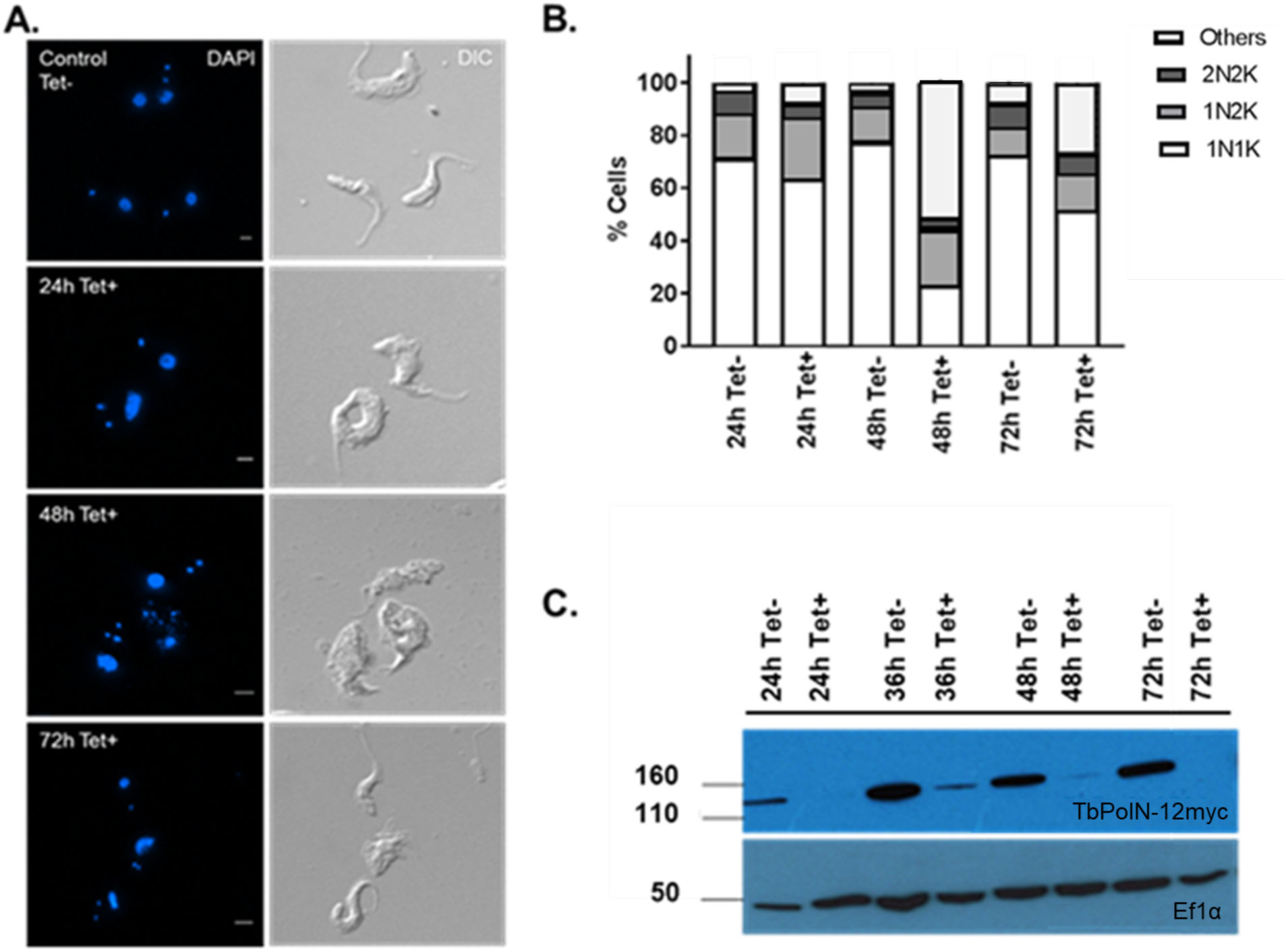
Analysis of the cell cycle after RNAi depletion of TbPolN. **A.** Cell cycle analysis by DAPI staining after 24, 48 and 72 hrs growth in the absence (Tet-) or presence (Tet+) of tetracycline RNAi induction. Immunofluorescence analysis allowing the visualization of the DNA content, and the cell body was visualised by differential interference contrast microscopy (DIC). **B**. Classification of Tet + and Tet-cells as 1N1K (G1/S), 1N2K (G2/M) and 2N2K (mitosis) cells at the time points shown; ‘others’ denotes cells that deviate from these classifications. Values depict the mean of each classification in the total population in two biological replicates. More than 200 cells were counted at each time point and in each experiment. **C.** Western blot analysis using anti-myc antiserum to evaluate abundance of TbPolN-12myc after 24, 48 and 72 hrs growth in Tet- and Tet+ cultures; detection of EF1a provides a loading control.

### TbPolN loss is associated with aberrant cell morphology

As detailed above, RNAi depletion of TbPolN resulted in significant changes in cell morphology, especially 48 hrs post induction. In order to evaluate these observations further, transmission electron microscopy (TEM) and scanning electron microscopy (SEM) were employed **(Fig.5).** TEM was performed in order to examine the ultrastructure of TbPolN RNAi depleted cells relative to uninduced controls, facilitating the visualization of the nucleus, kinetoplast and flagellar pocket. SEM allowed visualization of the external architecture of BSF cells. Upon depletion of TbPolN, clear alterations were observed in the internal morphology of the cell. Unlike in uninduced cells, the kinetoplast could not clearly be detected in the images of TbPolN RNAi depleted cells, even though DAPI staining suggested this DNA was present. However, cells with multiple flagellar pockets were seen. Since in *T. brucei* the kinetoplast is connected to the flagellum basal body [66], the number of kinetoplasts is normally directly related with the number of flagellar pockets, perhaps suggesting that DAPI imaging did indeed detect multiple kinetoplasts after depletion of TbPolN. The SEM images support this observation and reinforce the suggestion of aberrant cell division after TbPolN loss, since multiple flagella were readily detected in single cells. RNAi induced cells also presented irregular nuclei. Though this phenotype was not detected in all cells, it is consistent with aberrant nuclear staining with DAPI and suggests problems associated with replication or division of the nucleus. Finally, swollen ‘vacuole-like’ structures were found in a few cells. These structures seemed to be lacking any content and how they arose is unclear

**Figure 5.**
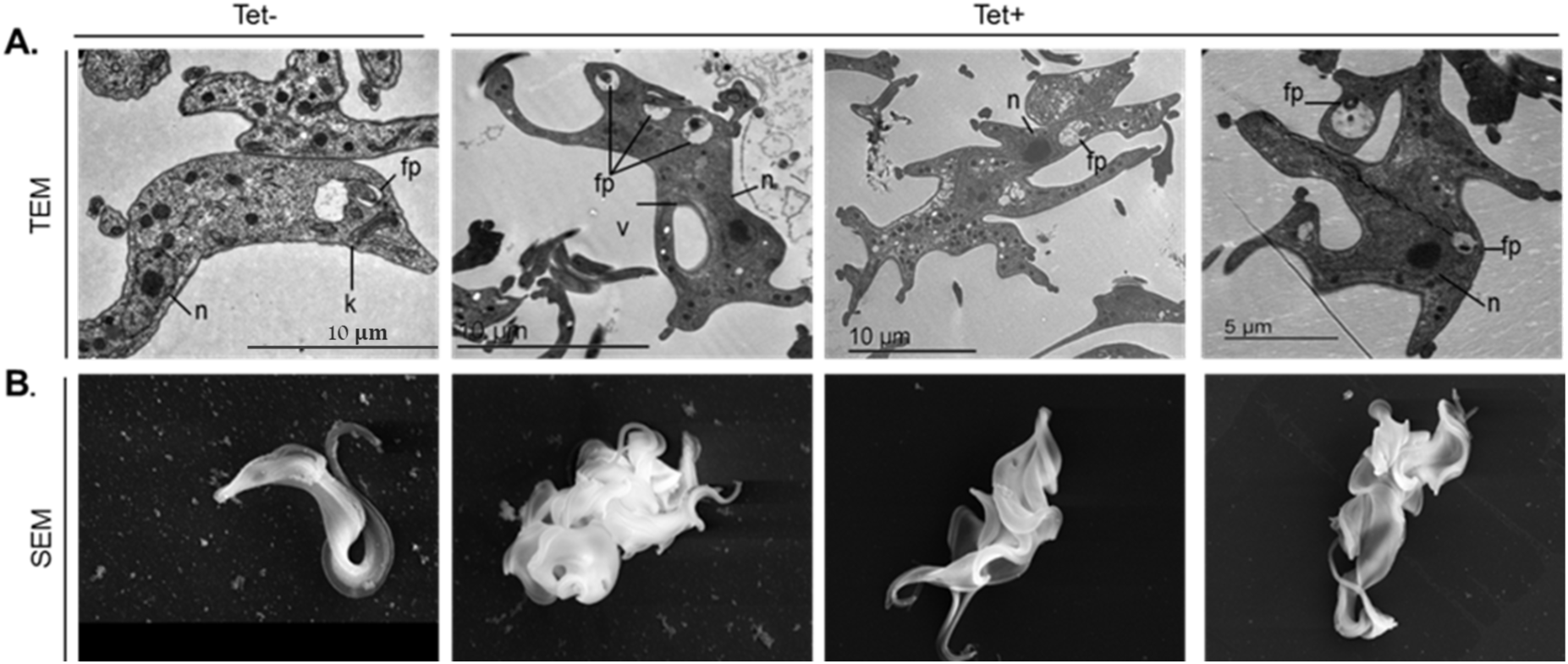
TbPolN loss is associated with aberrant cell morphology. **A.** Transmission electron microscopy images of *T. brucei* bloodstream forms before and after RNAi induction. The first panel shows a cell with a regular architecture, before the depletion of TbPolN (Tet-). Next three panels show representative images of aberrant cells 48 hrs post RNAi induction (Tet+), presenting abnormal internal architecture. Important features of the cells are indicated in black (n-nucleus; fp-flagellar pocket; and v-vacuole). **B.** Scanning electron microscopy images of *T. brucei* cells. The left panel shows a BSF *T. brucei* cell before RNAi induction (Tet-). The other three panels show representative images of aberrant cells 48 hrs post RNAi induction of TbPolN (Tet+).

### Loss of TbPolN results in nuclear damage in some cells

In *T. brucei* and *Leishmania* it has been demonstrated that Thr130 phosphorylation of histone H2A occurs in the presence of diverse types of damage [62] and in gene mutants [31];[67];[15], indicating this H2A modification is the kinetoplastid variant of serine phosphorylation of either H2A or the H2Ax variant in other eukaryotes. To assess the levels and distribution of γH2A after TbPolN RNAi, both immunofluorescence and western blotting were performed (Fig.6). Western analysis revealed that expression of γH2A was greater at both 24 and 48 hrs after TbPolN RNAi relative to uninduced cells. Immunofluorescence showed that a small amount of nuclear γH2A signal was seen in some uninduced cells, consistent with previous studies. After RNAi, only a modest increase in γH2A signal was seen in some cells after 24 hrs, whereas a substantially stronger signal was seen 48 hrs post RNAi induction (Fig.6, Fig.S5). Nevertheless, even 48 hrs after RNAi not every cell showed clear γH2A signal, and the level of nuclear anti-γH2A staining varied substantially between cells (Fig.S5), indicating the levels of nuclear damage resulting from loss of TbPolN were not uniform across the population. 72 hrs after RNAi γH2A signal decreased relative to the 48 hrs (Fig.6), mirroring the DAPI analysis of cell cycle changes (Fig.4).

**Figure 6.**
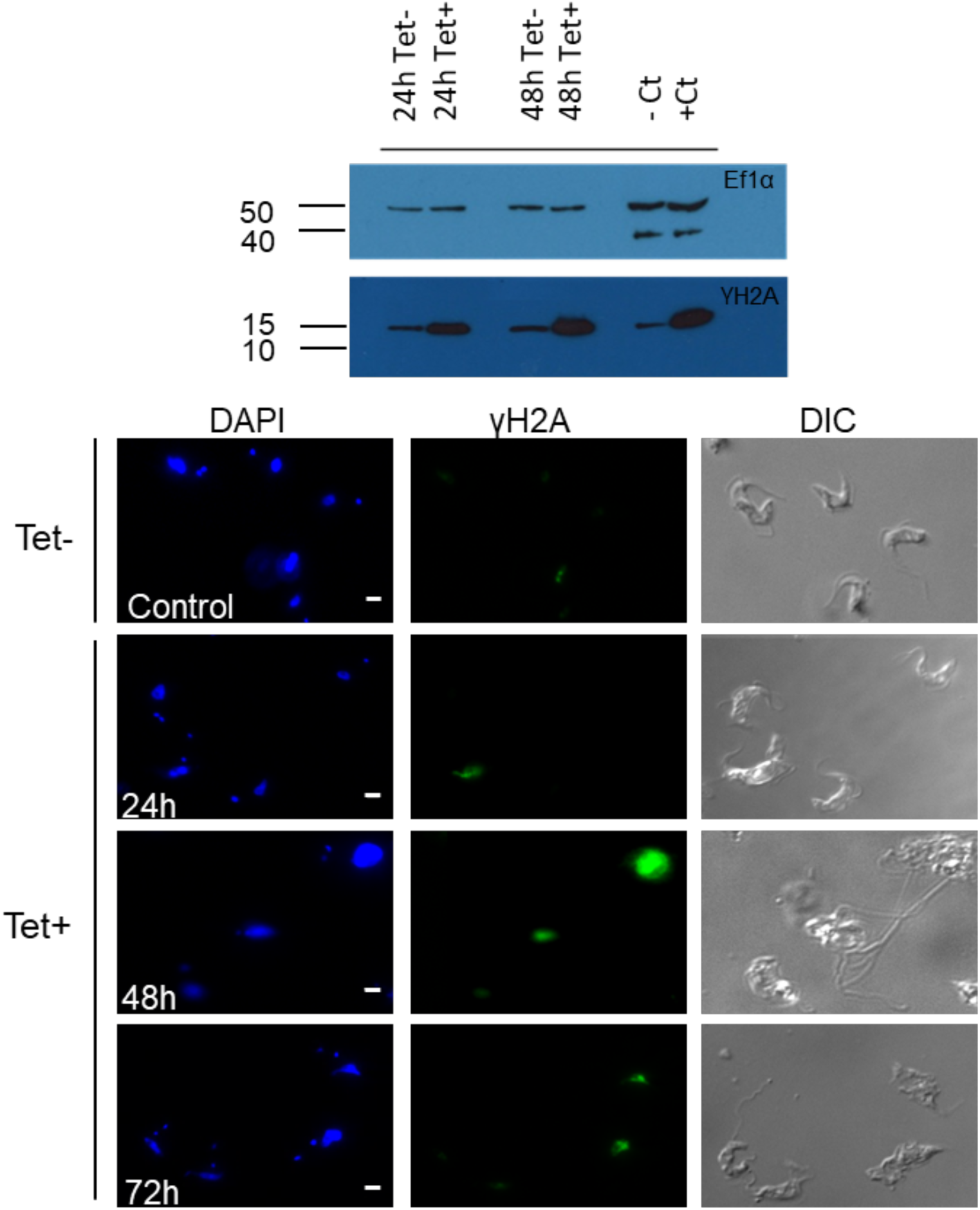
Increase in *T. brucei* γH2A after RNAi of TbPolN. **A.** Western blot showing γH2A levels after RNAi depletion of TbPolN relative to uninduced cells. Signal is shown after 24 hrs and 48 hrs of growth with (Tet+) or without (Tet-) RNAi induction, and detected with anti-γH2A antiserum. WT cells in the absence and presence of phleomycin (2.5 µg ml^−1^) were used as negative (-Ct) and positive (+Ct) controls, respectively. Anti-Ef1α antiserum was used as a loading control. Size markers (kDa) are shown. **B.** Immunolocalisation of *T. brucei* γH2A: left panels show DAPI staining of the nucleus and kinetoplast, the middle panels shows γH2A detected with anti-γH2A antiserum, and the right panel show the cell outline by DIC. Representative images are shown of cells 24 hrs, 48 hrs and 72 hrs after RNAi induction, or without induction (Tet-). Scale bar represents 2 µm.

### Depletion of TbPolN affects nuclear DNA replication

The above data reveal that nuclear damage accumulation, cell cycle abnormalities and a decrease in cell proliferation are all the result of RNAi depletion of TbPolN. One explanation for this range of phenotypes might be a role of the putative TLS Pol during nuclear genome replication, with genome instability when the protein is lost. To assess a role for TbPolN during DNA replication the capacity of the cells to incorporate EdU after TbPolN RNAi was assessed (Fig.7). The cells used in this assay were not synchronized and so were incubated with EdU for 4 hrs, on the basis that most cells should have undergone at least some DNA synthesis, assuming a nuclear S phase duration of around 1.5 hrs [68]. After 36 hrs and 48 hrs without RNAi, >90% of cells displayed nuclear EdU signal (Fig.S5). To allow comparison with EdU uptake after RNAi, the proportion of Tet-induced EdU positive cells at each time point was calculated relative to the uninduced cells at the same time point. A decrease in the number of EdU stained cells (22%) was detected 36 hrs after RNAi induction, and this became more pronounced after 48 hrs (when 45% of the population failed to incorporate EdU). These data suggest that DNA replication is impaired after TbPolN loss and concomitant with the emergence of cell cycle defects and accumulation of DNA damage.

**Figure 7.**
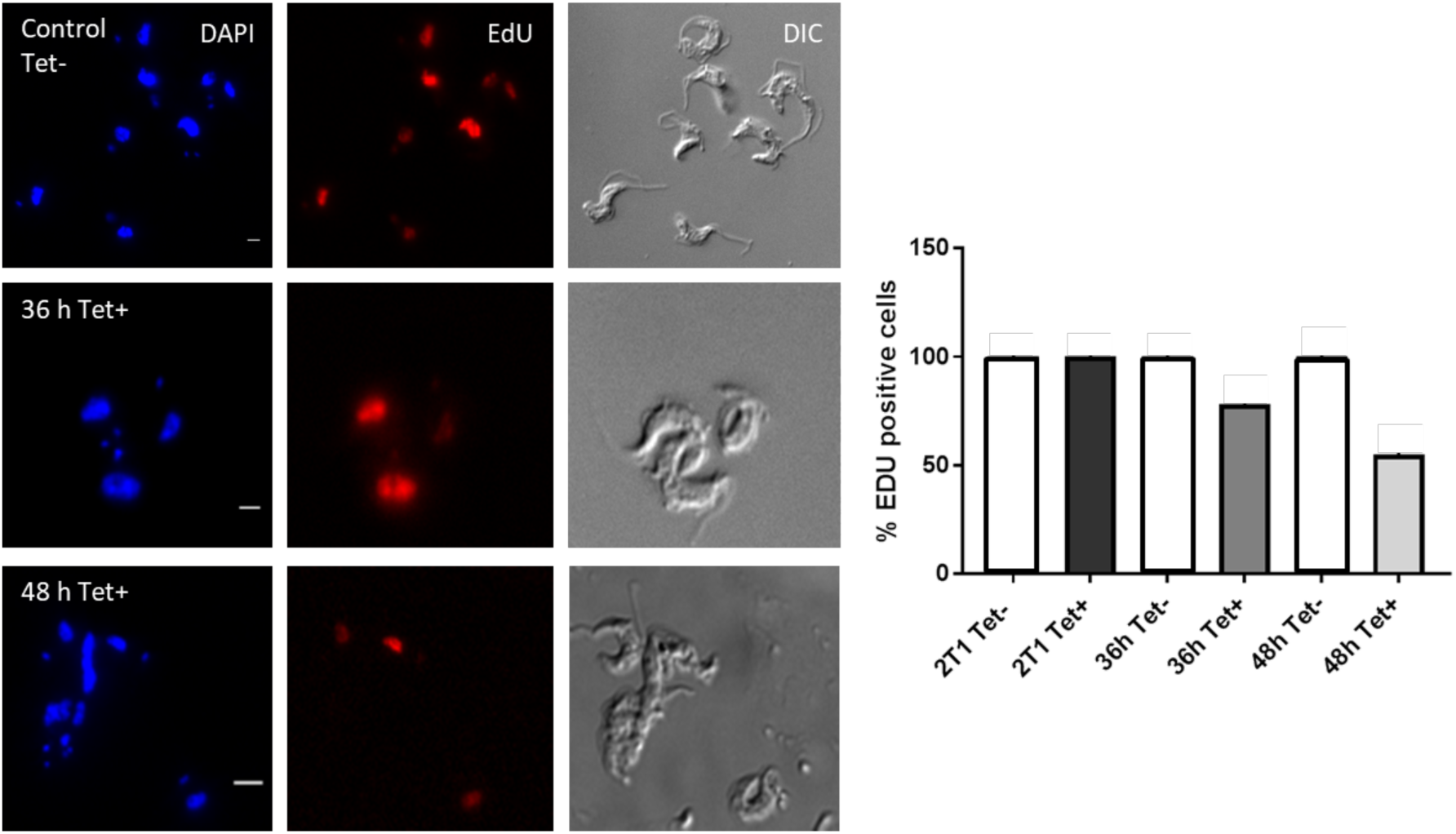
EdU labelling suggests impaired nuclear DNA replication after loss of TbPolN. **A.** Left panels show DAPI staining of the nucleus and kinetoplast, middle panels show EdU signal, and right panels show the cell outline by DIC. Representative images are shown of cells without RNAi induction (Control, Tet-) and after 36 or 48 hrs of growth after RNAi induction (Tet+). **B.** EdU positive cells without (Tet-) or with (Tet+) tetracycline were quantified after 36 hrs and 48 hrs of growth; the graph depicts the percentage of the total population with EdU signal, comparing the TbPolN RNAi induced cells at each time point with the same cells grown without Tet (set at 100%); parental 2T1 cells treated in the same way and grown for 48 hrs are also shown. A minimum of 100 cells were analysed per time point and bars represent mean values of two biological repetitions.

### Accumulation of DNA damage after TbPolN loss has a complex correlation with DNA replication

To ask if the accumulation of nuclear γH2A and impairment of DNA replication after TbPolN loss are linked, RNAi cells were grown for 48 hrs with tetracycline, stained for EdU uptake and simultaneously evaluated for γH2A expression by immunofluorescence (Fig.8). In the uninduced population, where few cells displayed γH2A signal, most had taken up EdU, meaning most (but not all) were negative for the former marker and positive for the latter. In contrast, many cells could be detected after TbPolN RNAi that did not display EdU signal but had γH2A signal (Fig.8, Fig.S5); indeed, this was commonly observed in aberrant cells. In addition, some cells without γH2A could be detected that had incorporated EdU (Fig.8, Fig.S5). Though these patterns may be consistent with stalling of nuclear DNA replication in those TbPolN RNAi cells that accumulate nuclear damage, cells were also detected, albeit less frequently, which were positive for both EdU and γH2A (Fig.8). These latter cells suggest that a simple cause and effect, such as genome damage resulting from TbPolN loss leading to impairment of DNA replication and growth, or vice versa, is not clear.

**Figure 8.**
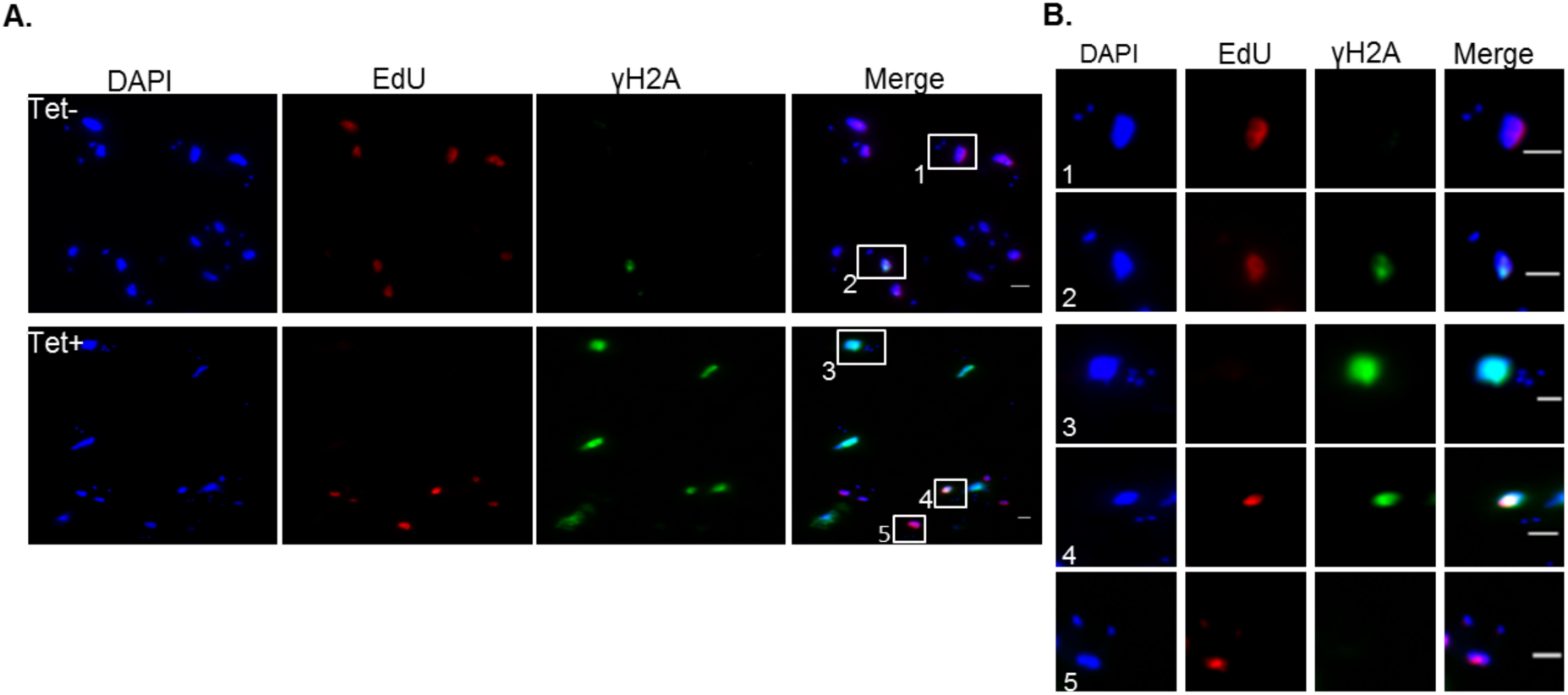
Immunofluorescence of both γH2A and EdU after TbPolN RNAi. **A.** Cells are shown after 48 hrs growth in the absence (Tet-) or presence (Tet+) of tetracycline RNAi induction. The first panels (left to right) show DAPI staining of the nucleus and kinetoplast, second panels show γH2A signal detected with anti-γH2A antiserum, third panels show EdU signal, and the fourth panels show a merge of the γH2A and EdU signals. **B.** Larger images of boxed areas show various patterns of the two nuclear signals seen in uninduced (Tet-) and induced (Tet+) cells: (1) cells showing EdU incorporation and lacking γH2A signal; (2) cells showing overlapping EdU and γH2A signal; (3) cells lacking EdU signal but showing accumulation of damage; (4) cells with overlapping γH2A and EdU signal; and (5) cells incorporating EdU and not showing γH2A signal. Scale bar 2 µm.

### Loss of TbPolN results in aberrant chromosome segregation in a subset of cells

Although PolN mutation has not been associated with chromosomal abnormalities in other eukaryotes, the severity of the phenotypes observed after RNAi depletion of TbPolN prompted us to test for loss of chromosome integrity. In order to do so, a telomere-fluorescence *in situ* hybridisation (Telo-FISH) assay was performed on TbPolN RNAi cells after 24 hrs and 48 hrs with and without RNAi induction (Fig.9). Because mini-chromosomes are more abundant than megabase-chromosomes, it is likely that most of the signal seen using a telomere probe corresponds to the former molecules. As seen in previous work [69], in uninduced RNAi cells, telomere signal was seen diffusely around the nucleus during interphase, with a stronger signal towards the nuclear envelope [70]. During metaphase the telomeres were found in the centre of the nucleus, and during anaphase the telomeres localised towards the nuclear poles [70];[69]. From 24 hrs after RNAi induction, corresponding with the emergence of growth arrest, cell cycle abnormalities and DNA damage, cells with aberrant telomere signals could be seen (Fig.9, further examples Fig.S6). Most interphase cells appeared comparable with the controls, whereas the typical telomere distribution during metaphase was often absent after RNAi, with cells detected that lacked the expected signal at the equator of the nucleus. During anaphase the ordered segregation of the telomeres into each of the two progeny cells could also be seen to be compromised, with unequal distribution of signal in the nuclei. Finally, telomere distribution was notably unusual in aberrant cells, which were mainly seen after 48 hrs of RNAi and whose N-K content meant their stage in the cell cycle was unclear. Taken together, these data may be explained by inaccurate chromosome segregation events during mitosis in some cells after loss of TbPolN, which appears concomitant with the emergence of cells with abnormal N-K ratios and consistent with loss of the putative TLS Pol not impeding mitosis.

**Figure 9.**
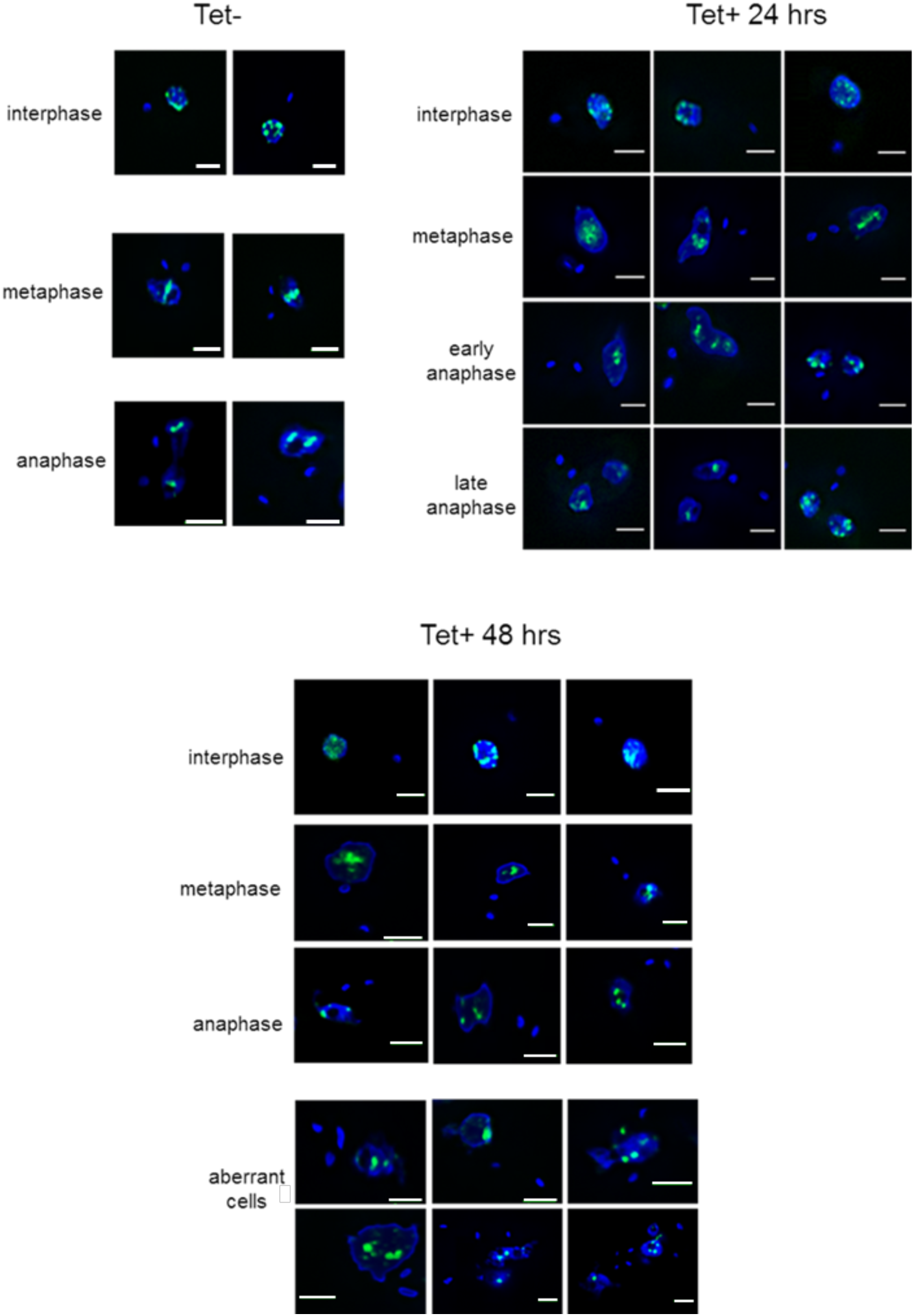
Loss of TbPolN affects chromosome segregation during cell division. Fluorescence *in situ* hybridization using telomeres as a probe (green) 24 or 48 hrs after tetracycline induction of RNAi against TbPolN (Tet+), or in the absence of tetracycline RNAi induction (Tet-). Representative images are shown of cells in interphase, metaphase and anaphase (separated into early or late stages for Tet+ 24 hrs cells), as well as aberrant cells whose cell cycle phase is unclear. DNA is stained with DAPI; scale bars: 2 µm.

### Loss of TbPolN leads to increased expression of silent VSG genes

Given our previous demonstration that TbPolN binds to chromatin containing telomeric repeats [16], and the above evidence that loss of TbPolN can affect telomere distribution during nuclear division, we next asked if loss of TbPolN affects the expression of VSGs from telomeric bloodstream VSG expression sites (Fig.10). Whole cell lysates were prepared at different time points after induction of RNAi and analysed with antibodies specific for VSG-2, VSG-3 or VSG-13. The parental cell line used to derive the RNAi cells used here encodes only VSG-2 (Fig.10), and the same predominant expression of this VSG was seen at time 0, with levels of VSG-2 only seen to decrease slightly 72 hrs post induction (Fig.10A). In contrast, RNAi induction led to a clear increase in signal for VSG-3 or VSG-13, which are normally transcriptionally repressed in this *T. brucei* strain and were barely detected at time 0. Intriguingly, the extent of the expression of both normally silent VSGs was highest 48 hrs post induction, consistent with the dynamics of the cell cycle and DNA damage effects described above. The same effects of increased expression of normally silent VSGs were clearly detectable in another, independent RNAi cell line (Fig.S7), which suggests either derepression of silent VSG genes or increased switching induced by TbPolN depletion.

**Figure 10.**
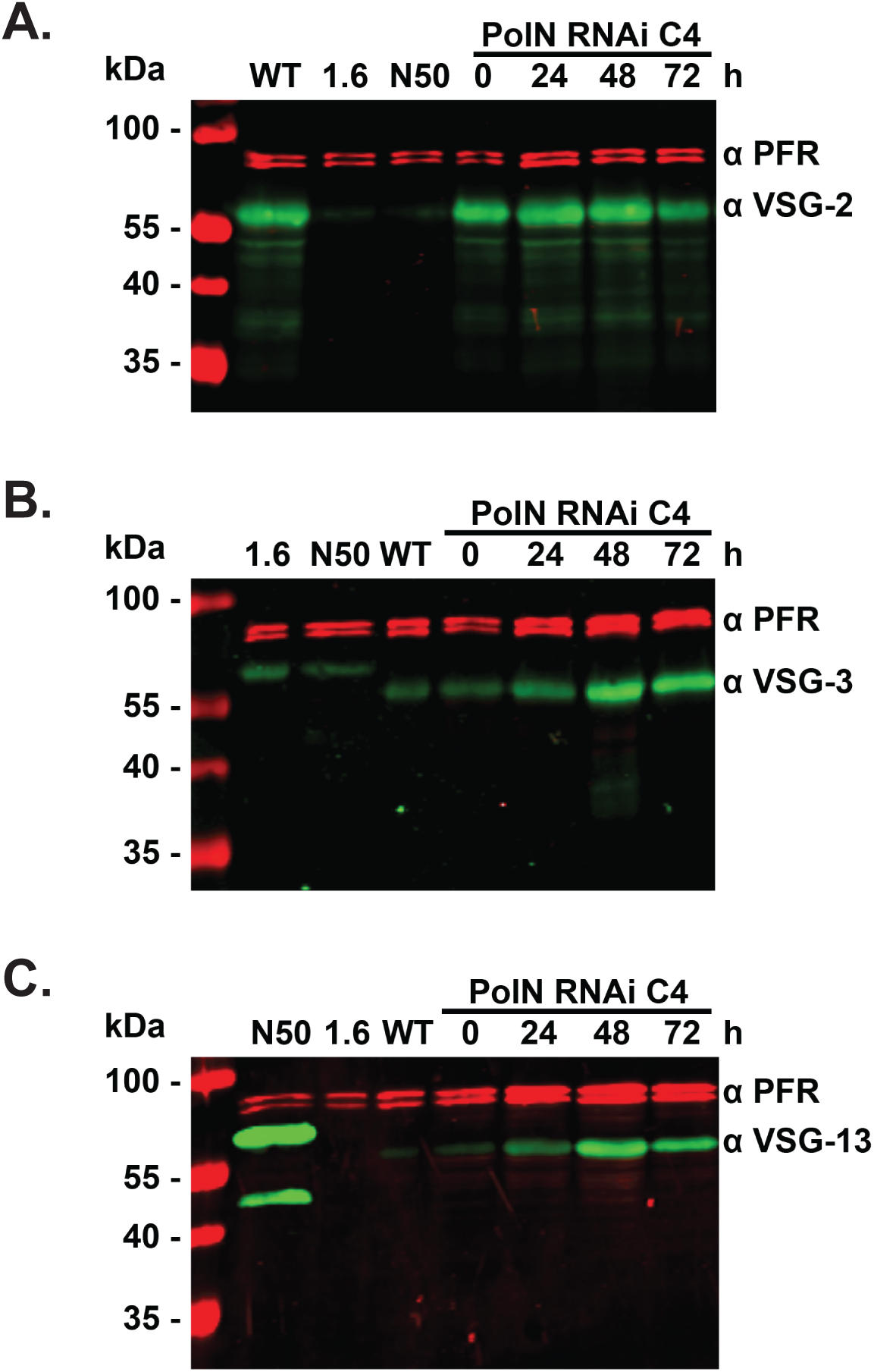
Increased expression of silent VSGs after RNAi depletion of TbPolN. Whole cell lysates were prepared at different time points after induction of RNAi against PolN in one clone (C4) and probed, after blotting, with antibodies specific for VSG-2 (**A**), VSG-3 (**B**) and VSG-13 (**C**). Antiserum against paraflagellar rod protein (PFR) was used as loading control. In addition, VSG levels were evaluated in a number of control cell lines: MiTat1.2 cells (WT) express VSG-2, whereas M1.6 (MITat1.6) express VSG-6, and N50 express VSG-13.

### Mass spectrometry analysis shows increased expression of VSGs from throughout the silent archive after PolN depletion

To attempt to distinguish between loss of transcriptional control and increased switching, we sought to determine the range of VSGs that become expressed in TbPolN-depleted parasites (Fig.11). To do so, we used a recently published protocol [71] to isolate and characterise the abundance of VSGs from two independent TbPolN RNAi cell lines 48 and 72 hrs after RNAi induction. Parental 2T1 cells were analysed as a negative control and, like the uninduced RNAi cells, mainly expressed VSG-2 and showed only very minor traces of VSG-8 expression. In contrast, we could detect several VSGs after depletion of TbPolN. In all samples the abundance of VSG-8, as well as VSGs from several further BES, including VSG-6, VSG-17 and VSG-11, increased (Fig11 and Table 1). In addition, substantial increased levels were detected of VSG-653, which is present in a metacyclic VSG expression site (MES) and normally only expressed in the tsetse fly. Finally, we could identify increased abundance of several VSG, such as VSG-636 and VSG-336, which are located in sub-telomeric regions of chromosomes 8 and 9 and not within a BES or MES. Taken together, these data suggest that increased VSG expression after loss of TbPolN may not simply result from derepression of normally silent telomeric VSG genes in the BES and, perhaps, the MES. The observation that VSGs from the silent, subtelomeric arrays are also found to increase may suggest that loss of TbPolN also increases recombination events between subtelomeric VSG genes and the BES.

**Figure 11.**
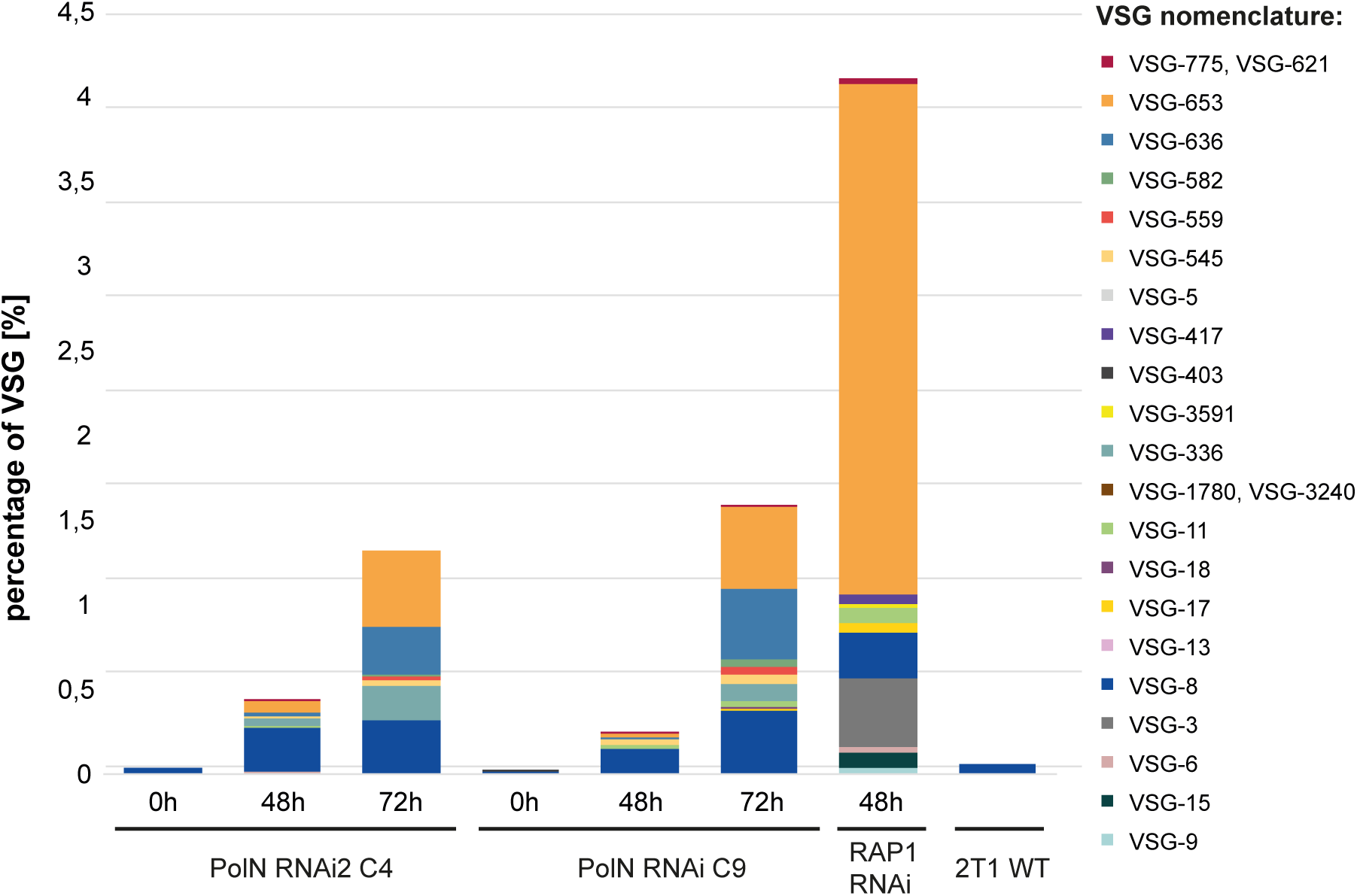
Mass spectrometry reveals increased expression of multiple silent VSGs after TbPolN loss. The graph shows VSGs identified by mass spectrometry 0, 48 and 72 hrs after induction of RNAi against TbPolN. Data is shown for two RNAi clones (C4 and C9), VSG identifiers are coloured, and VSG abundance in the different time points is shown as percentage representation in the samples. VSG-2 is not shown as it displayed an abundance of 95-98 % in each sample. The parental 2T1 (WT) cell line was analysed as negative control, and VSGs expressed after 48 hrs of TbRAP1 RNAi, which is known to result in de-repression of inactive VSG BESs is shown for comparison.

**Table 1.** List of identified VSGs in TbPolN depleted trypanosomes by mass spectrometry. VSG-2 is not listed, because of its abundance of 95 – 98 % in each sample. VSGs were identified in two independent clones, except for VSGs highlighted by light grey boxes, which were only found in TbPolN RNAi C9 cell line. MBC: Megabase chromosome; IC: Intermediate chromosome; STR: sub-telomeric region; BES: bloodstream

| VSG | Localisation in ES | length of ES | Localisation in the Genome |
| --- | --- | --- | --- |
| VSG-6 | BES-3 | approx. 47 kb | MBC 4 |
| VSG-8 | BES-12 | approx. 42 kb | MBC 2 |
| VSG-17 | BES-13 | approx. 52 kb | IC |
| VSG-18 | BES-5 | approx. 42 kb | MBC 5 |
| VSG-11 | BES-15 | approx. 58 kb | MBC 3 |
| VSG-1780,<br>VSG-3240 |  |  | unknown |
| VSG-336 |  |  | 3'-STR MBC 8 |
| VSG-403 |  |  | 3'-STR MBC 1 |
| VSG-5 |  |  | 3'-STR MBC 8 |
| VSG-545 |  |  | 3'-STR MBC 8 |
| VSG-559 |  |  | 5'-STR MBC 9 |
| VSG-582 | atypical MES |  | 3'-STR MBC 5 |
| VSG-636 |  |  | 5'-STR MBC 9 |
| VSG-653 | MES |  | 5'-STR MBC 11 |
| VSG-775,<br>VSG-621 |  |  | unknown,<br>3'-STR MBC 10 |

## Discussion

Translesion DNA Pols, in addition to allowing cells to grow and survive in the presence of hard to repair genome lesions, are increasingly implicated in wider genome functions, including in various repair pathways [72];[73], telomere homoeostasis [74];[75] and replication initiation [45]. In this work we have explored the functions of PolN in *T. brucei* and document that this putative TLS Pol, in addition to having important roles in genome maintenance, acts in antigenic variation, a strategy for immune evasion found in a diverse array of bacterial, fungal and protozoan pathogens.

Relatively little work has explored the functions of TLS PolN in other eukaryotes, but null mutants of the gene are viable in mice [51]. Loss of PolN has been described to affect meiotic recombination, with conflicting evidence for the protein having a role in tackling cross-link DNA damage [51];[50]. One reason that has been proposed for this limited functionality is that PolN may have arisen recently in evolution, having only been described to date in metazoa [51]. This limited phyletic distribution contrasts with PolQ, which is found widely in eukaryotes and shares DNA Pol domain homology with PolN. Here we show that PolN, but not PolQ, is present in *T. brucei* and plays a more critical role in the parasite that has been described in mice, since RNAi-mediated loss of TbPolN results in substantial growth impairment that is concomitant with cell cycle defects, accumulation of nuclear damage, reduced DNA synthesis, and chromosome defects. The presence of PolN in *T. brucei* and related kinetoplastids may suggest that this TLS Pol is more widespread than has previously suggested. However, our naming of the protein as PolN reflects the fact that the polypeptide encodes only a PolQ-like DNA Pol domain, and lacks an SNF2 helicase domain (which is present but encoded from an unlinked gene whose function has not been examined in *T. brucei*). In *Leishmania*, previous researchers called the protein encoded by the syntenic and orthologous gene PolQ [60], which we suggest is formally incorrect, due to the lack of linked Pol and helicase domains that define PolQ. Nonetheless, the lack of a dual function PolQ in kinetoplastids may suggest that this eukaryotic lineage evolved separate PolN-like and HelQ-like proteins from a PolQ ancestor relatively recently, and the more severe effects of loss of PolN in *T. brucei* than in mice and human cells reflect retention of some PolQ-like roles. Why such a putative gene fission may have evolved in kinetoplastids is unclear, and merits further testing. For instance, we do not know if TbPolN and the putative TbHelQ function together, as has been suggested for PolN and HelQ (Hel308) in human cells [50].

Our analysis of TbPolN represents just the second functional examination of a TLS Pol in *T. brucei* and, perhaps surprisingly, both proteins are critical for growth of the parasite in culture. Previously, Rudd et al examined two Prim-Pol proteins in *T. brucei*, called TbPPL1 and TbPPL2 [15]. Loss of TbPPL2 by RNAi, like loss of TbPolN, dramatically impairs cell growth and leads to accumulation of damage. However, several lines of evidence suggest that the proteins perform distinct functions. First, TbPolN-12myc displays a striking localisation mainly in two peripheral regions of the nucleus, whereas no such discrete accumulation of TbPPL2 was described. Second, though loss of both Pols results in increased γH2A signal in the nucleus, indicating accumulation of damage, the modified histone is seen as multiple foci after TbPPL2 RNAi, whereas after TbPolN loss γH2A signal is not clearly focal and displays considerable variation between cells. Third, though loss of TbPolN caused redistribution of the Pol in the nucleoplasm after alkylation damage by MMS, we found limited evidence for recruitment of the protein to the resulting γH2A foci, unlike the pronounced overlap of γH2A and TbPPL2 seen after MMS exposure. Finally, whereas loss of TbPPL2 resulted in a striking stall of cell division at G2/M, loss of TbPolN appears instead to have a less severe impediment to cell cycle progression, since cells with aberrant DNA content arise after RNAi that are indicative of inaccurate genome segregation after mitosis. It is conceivable that this difference in cell cycle response explains why loss of TbPPL2 appears to have a more severe effect on cell growth than loss of TbPolN. However, it is less clear why the cell cycle and DNA damage defects that arise after loss of TbPolN peak 48 hrs after RNAi and then diminish; whether this indicates adaptation of the cells, such as by increased use of an alternative TLS Pol, is unclear. Despite such disparate activities, the fact that loss of either protein results in such pronounced loss of cell viability is striking, since it indicates endogenous genome features in *T. brucei* that must be tackled by both TLS Pols for effective genome transmission during cell division. What such endogenous genome features might be is unknown. TbPPL2 has been shown to allow replication bypass of base cross-links generated by UV and 8-oxoG formed by oxidation, but is not capable of bypassing 3dMeA or an abasic site [15]. LiPolN (LiPolQ), the likely *Leishmania* orthologue of TbPolN, shares some but not all of these catalytic activities: like TbPPL2 it can bypass 8-oxoG, but is also capable of crossing an abasic site [60]. Whether any of these activities underlie the important, endogenous roles played by the TLS Pols is unknown, and we cannot exclude the possibility that the defects seen after loss of either protein reflect a shared and hitherto undescribed role. Notably, previous analysis did not ask if loss of TbPPL2 affects VSG expression, as we describe after RNAi of TbPolN.

Our previous work revealed association of TbPolN with the *T. brucei* telomere [16], but what role it may play was not tested. To date, PolN has not been associated with telomere homeostasis in any eukaryote, though TLS Pol eta activity has been shown to contribute to alternative lengthening of telomeres in mammalian cells [74], where it may contribute to recombination [76]. No such activity has been described for PolN in any organism, and analysis of the potential roles of kinetoplastid TLS Pols in recombination is so far limited to *T. cruzi* [77];[78], without examining the telomere. Our data show that loss of TbPolN results in changes in expression of VSG genes that are found adjacent to the telomere with evidence for increased expression of VSG in BES and MES, as well as subtelomeric array VSGs. We cannot say if these alterations are due to an impact of TbPolN loss on telomere maintenance, but the effects on VSG expression controls have some similarity with those seen after loss of *T. brucei* shelterin components, including TbRAP1, TbTRF and TbTIF2. All three of these telomeric factors are essential for trypanosome viability. However, while TbTRF and TbTIF2 only play minor roles in VSG silencing, TbRAP1 is critical for BES-linked VSG suppression [23]. Depletion of TbRAP1 leads to derepression of all silent BESs, with the most prominent effect at telomere-proximal regions. In addition, a recent study [32] revealed that TbRAP1 loss leads to increased read-through into the telomere downstream of the active BES, resulting in higher levels of non-coding telomeric repeat-containing RNAs (TERRA) and telomeric RNA:DNA hybrids. In addition, more DSBs were detected at telomeres and subtelomeres of active and silent BESs, which increased the VSG switching frequency by initiating VSG gene conversion. In contrast, depletion of TbTRF and TbTIF2 did not affect TbRAP1-mediated VSG silencing, but resulted in increased gene conversion-mediated VSG switching events [26], which appeared to rely on independent and overlapping mechanisms, since TbTIF2 knockdown leads to the accumulation of subtelomeric DSBs, an effect not seen after TbTRF depletion. This mechanistic overlap may arise because TbTIF2 stabilizes TbTRF protein levels by suppressing its degradation by the 26S proteasome [26]. In summary, although TbTRF, TbTIF2 and TbRAP1 are components of the same complex, only TbRAP1 is crucial for telomeric VSG silencing, suggesting that TbTIF2 and TbTRF act independently from TbRAP1, which is supported by yeast two-hybrid studies that reveal only a weak interaction between TbRAP1 and TbTRF [23], but a very strong interaction of TbTRF and TbTIF2 [25].

Given the above complexity in VSG switching functions of the three known telomere-associated factors in *T. brucei*, it is possible that TbPolN also contributes to telomere stability and its ablation leads to damage that undermines monoallelic VSG expression and/or leads to VSG recombination. Such a telomere-directed function would be consistent with our previous demonstration that TbPolN binds chromatin-associated telomeric repeats [16], and the observation here that RNAi against TbPolN alters telomere localisation. However, other explanations might also be considered. One alternative possibility is that TbPolN plays a direct role in VSG switching, such as during replication of the VSG BES or contributing to VSG recombination. For instance, TbPolN may play an important role in overcoming impediments to replication that are found in the VSG BES and, in its absence, damage accumulates and increases VSG switching. What such impediments might be are unclear, but RNA-DNA hybrids [31] and a modified base (J) [79] have been described in the VSG BES, as well as elsewhere in the genome [31];[80]. If TbPolN were to play a catalytic role in VSG recombination, it is unclear why its depletion would lead to the observed increases in silent VSG expression, rather than impeding changes in VSG expression. A wider explanation may be that loss of TbPolN affects genome replication, which then deregulates the controls that normally ensure monoallelic VSG expression. For instance, it is known that the active VSG BES, unlike all the silent BESs, is replicated early in S phase [27]. If S phase progression or the correct segregation of chromosomes were perturbed, it is conceivable that such targeted replication of only the active BES might breakdown, leading to changes in VSG transcription or recombination in all VSG BES. Such a replication-focused explanation may explain why similar effects on VSG expression are seen after depletion of the factors involved in DNA replication, including the Origin Recognition Complex [28];[29] and MCM-BP [30]. However, whether such effects are due to global problems with genome replication, or are due to VSG BES-focused roles is difficult to separate. However, in this regard, two observations are worth noting. First, RNAi depletion of TbPPL2 does not lead to detectable effects on VSG expression (data not shown), which may be because loss of this TLS Pol prevents transition from G2/M into mitosis [15], unlike the cell cycle perturbations seen after loss of TbPolN. Second, recent data have implicated HelQ of *T. cruzi* in DNA replication [56]; if such a role is conserved in *T. brucei*, and if trypanosomatid HelQ and PolN proteins work together, then a link between VSG switching and genome replication might be more firmly established.

## Acknowledgement

Work in the Butter and Janzen laboratories was supported by a joint DFG research grant (BU2996/13-1, JA1013/6-1).) Work in the McCulloch laboratory was supported by the BBSRC [BB/K006495/1, BB/M028909/1, BB/N016165/1 and a DTP studentship to E. Briggs] and by a SENESCYT (Secretaria Nacional de Educación Superior, Ciencia y Tecnología e Innovación) PhD scholarship to AL. The Wellcome Centre for Integrative Parasitology is supported by core funding from the Wellcome Trust [104111]

## Author contributions

R.M., F.B. and C.J.J. designed the study. A.Z.L., M.S., E.B., H.R. and L.L. performed the bioinformatics analysis, bio-imaging and cell biology analysis of the TbPolN-depleted parasites. K.L. and F.B. performed the mass spectrometry analysis. R.M., F.B. and C.J.J. wrote the manuscript. All authors read and approved the final manuscript.

## Supporting information legends

**Figure S1. TbPolN-12myc nuclear localisation in different cell cycle stages in the absence or presence of alkylation damage. A.** Western blot of whole cell extracts of wild-type cells (WT) and cells in which the TbPolN gene was translationally fused with 12 myc (TbPolN-12myc); the blot was probed with anti-myc and anti-EF1a antisera, and the identity of the proteins detected is indicated. **B., C.** Immunolocalisation of TbPolN-12myc in *T. brucei* BSF cells grown in culture without damage (-MMS, B) or after 18 hrs growth in the presence of 0.003% methyl methanosulphonate (+MMS, C). The left panels show DAPI staining of DNA, the middle panels show signal detected with a conjugated Alexa Fluor® 488 anti-myc antibody, and the right panels are merged images. Scale bar represents 2 µm.

**Figure S2. Co-localisation of TbPolN-12myc and γH2A.** Immunolocalisation of *T. brucei* γH2A and TbPolN-12myc: first (left to right) panels show DAPI staining of DNA, the second panels show TbPolN-12myc using an unconjugated anti-myc antibody, third panels shows γH2A signal, and the fourth panels are merged images of the three signals. Cells are shown after growth for 18 hrs in the presence of 0.0003% of MMS (+), or without exposure to MMS (-). Images were captured on a Delta Vision RT deconvolution microscope. Scale bars represent 2 µm.

**Figure S3. Propidium iodide staining of unfixed cells. A.** Parasites were stained with propidium iodide without previous fixation to evaluate the amount of dead cells after induction of RNAi: two different clones (C4 and C9) are shown at indicated time points after RNAi induction. Gates define populations that are alive (A) or dead (D). Parental cells and puromycin-treated parasites served as negative (living cells) and positive (dead cells) controls, respectively**. B.** Quantification of flow cytometry profiles. Gates are shown in Fig. S3A. (n=3, **:p<0.01, ***:p<0.001, unpaired t-test); DC denotes puromycin treated, and WT denotes untreated.

**Figure S4. Loss of *T. brucei* PolN causes a modest increase in sensitivity to two forms of DNA damage.** Growth of *T. brucei* cells is shown in the presence of increasing concentrations of MMS (A) or after exposure of increasing doses of UV radiation (B), in each case without induction of RNAi (left graphs) or after induction of RNAi (right graphs). Average cell density is shown at three time points and error bars denote standard deviation from three experiments.

**Figure S5. EdU and γH2A fluorescence intensity after TbPolN RNAi.** Fluorescence intensity of EdU and γH2A (gamma histone) signal was calculated in Image J as the corrected total cell fluorescence (CTCF), using a region of interest (ROI, 21×21 pixels) around the cell and subtracting background signal. Dots represent the signals obtained for each individual cell, which were either induced for RNAi (Tet+) for 48 hrs, or grown for the same length of time without induction (Tet-). A total of 50 cells were analysed for each group.

**Figure S6. Fluorescence *in situ* hybridization of telomeres 48 hrs after depletion of TbPolN.** Representative images of cells after fluorescence *in situ* hybridization using telomeres as a probe (green) and after 48 hrs growth with induction of RNAi against TbPolN. Top panels show telomere distribution in 1N1K cells, middle panels show telomere distribution in 1N2K cells, and bottom panels show telomere localisation in aberrant cells. DAPI is in blue; scale bar, 2 µm.

**Figure S7. Increased expression of silent VSGs after RNAi depletion of TbPolN.** Whole cell lysates were prepared at different time points after induction of RNAi against PolN in clone C9 and probed, after blotting, with antibodies specific for VSG-2 (A), VSG-3 (B) and VSG-13 (C). Antiserum against paraflagellar rod protein (PFR, red) was used as loading control. Control cell lines (left panel) were taken from Figure 10.

