## Supplemental figures for "Genome maintenance functions of *Trypanosoma brucei* DNA Polymerase N include telomere association and a role in antigenic variation"

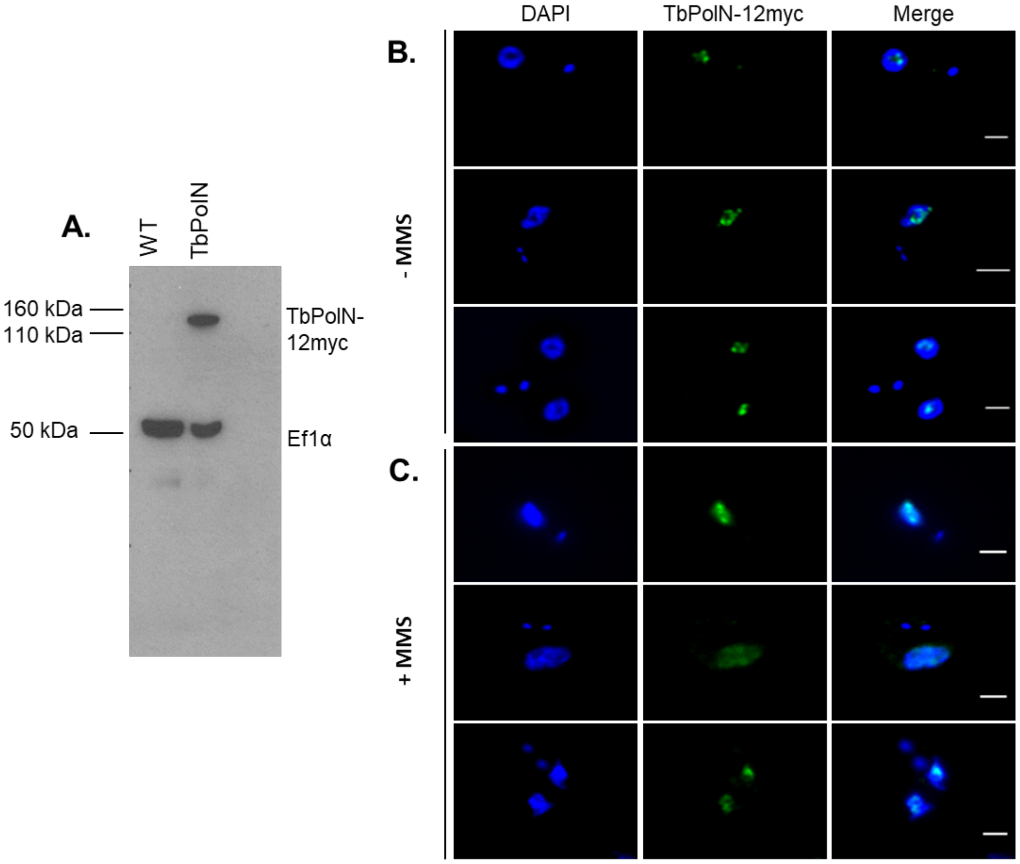


**Figure S1. TbPolN-12myc nuclear localisation in different cell cycle stages in the absence or presence of alkylation damage. A.** Western blot of whole cell extracts of wild-type cells (WT) and cells in which the TbPolN gene was translationally fused with 12 myc (TbPolN-12myc); the blot was probed with anti-myc and anti-EF1a antisera, and the identity of the proteins detected is indicated. **B., C.** Immunolocalisation of TbPolN-12myc in *T. brucei* BSF cells grown in culture without damage (-MMS, B) or after 18 hrs growth in the presence of 0.003% methyl methanosulphonate (+MMS, C). The left panels show DAPI staining of DNA, the middle panels show signal detected with a conjugated Alexa Fluor® 488 anti-myc antibody, and the right panels are merged images. Scale bar represents 2 μm.

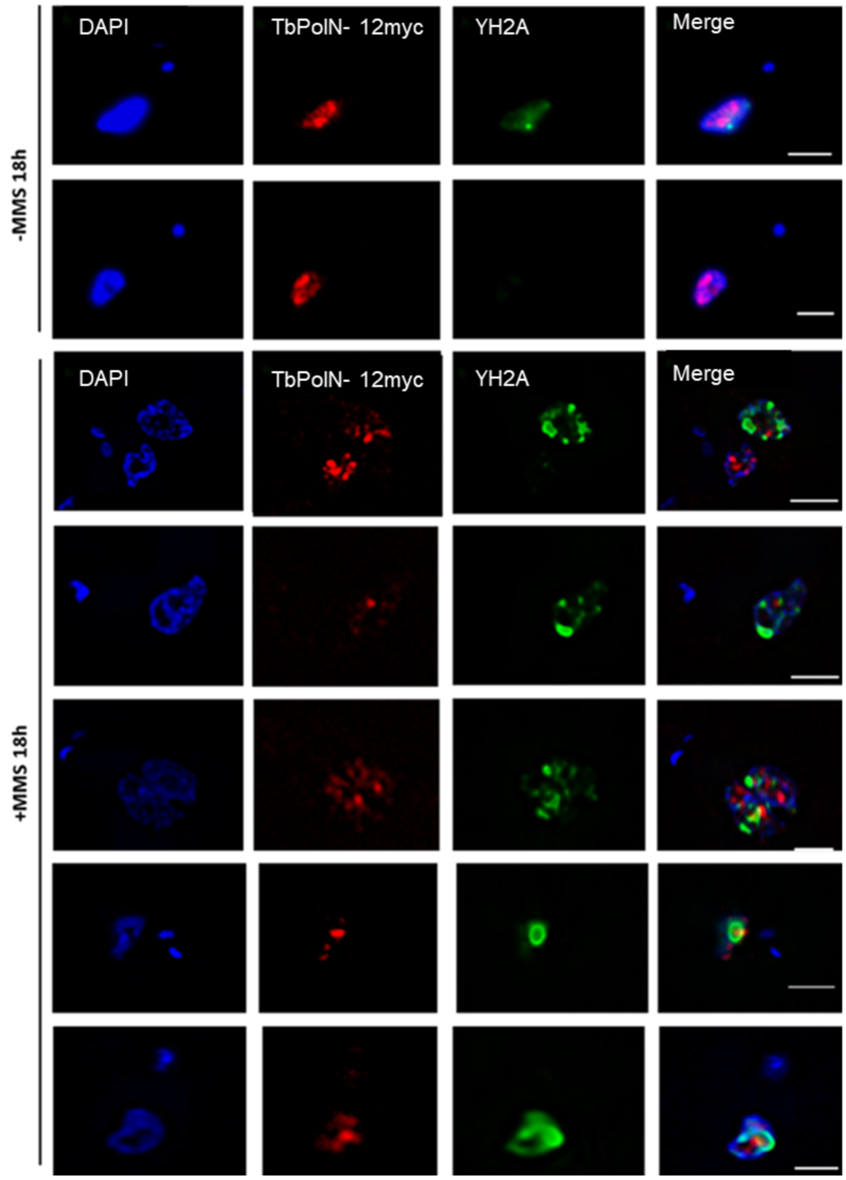


**Figure S2. Co-localisation of TbPolN-12myc and γH2A.** Immunolocalisation of *T. brucei* γH2A and TbPolN-12myc: first (left to right) panels show DAPI staining of DNA, the second panels show TbPolN-12myc using an unconjugated anti-myc antibody, third panels shows γH2A signal, and the fourth panels are merged images of the three signals. Cells are shown after growth for 18 hrs in the presence of 0.0003% of MMS (+), or without exposure to MMS (-). Images were captured on a Delta Vision RT deconvolution microscope. Scale bars represent 2 μm.

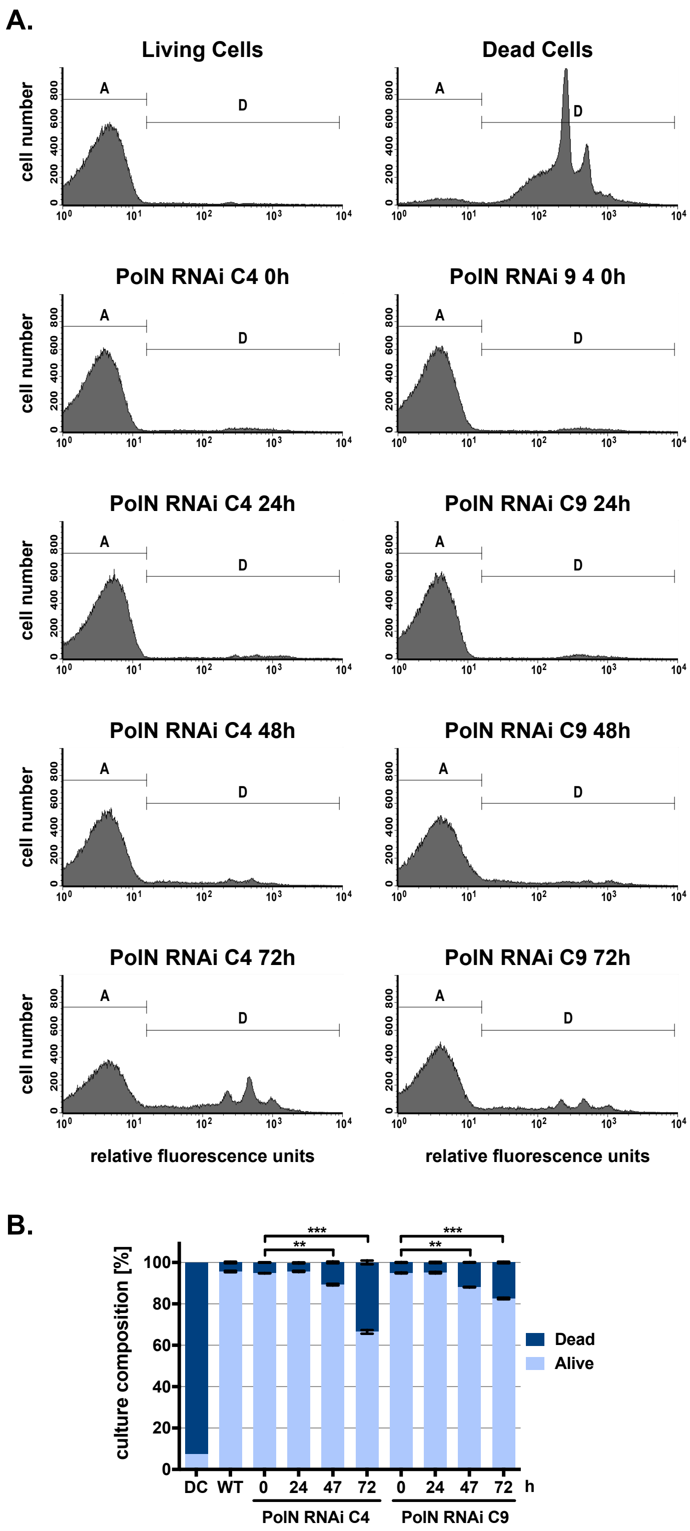


**Figure S3. Propidium iodide staining of unfixed cells. A.** Parasites were stained with propidium iodide without previous fixation to evaluate the amount of dead cells after induction of RNAi: two different clones (C4 and C9) are shown at indicated time points after RNAi induction. Gates define populations that are alive (A) or dead (D). Parental cells and puromycin-treated parasites served as negative (living cells) and positive (dead cells) controls, respectively**. B.** Quantification of flow cytometry profiles. Gates are shown in Fig. S3A. (n=3, **:p<0.01, ***:p<0.001, unpaired t-test); DC denotes puromycin treated, and WT denotes untreated.

**A**

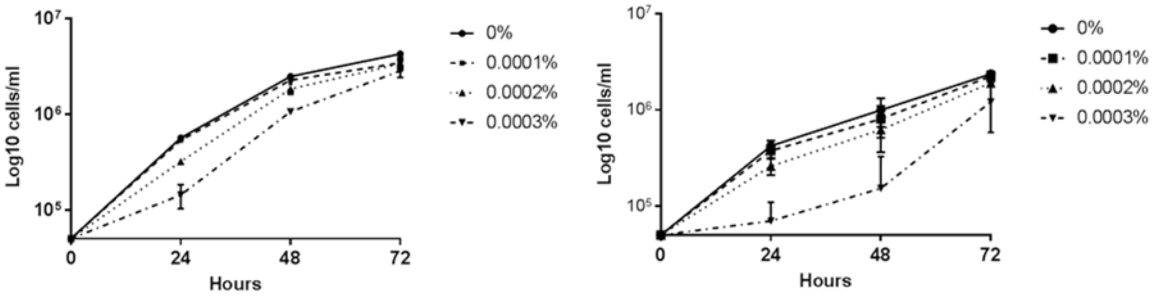


**B**

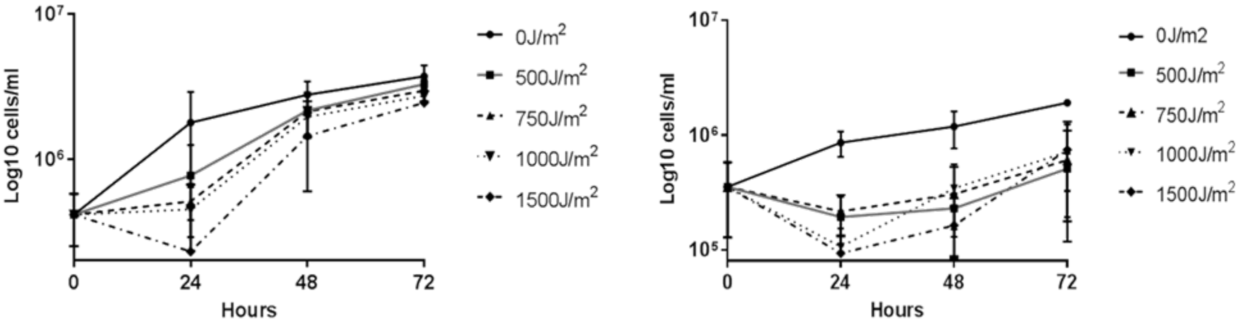


**Figure S4. Loss of *T. brucei* PolN causes a modest increase in sensitivity to two forms of DNA damage.** Growth of *T. brucei* cells is shown in the presence of increasing concentrations of MMS (A) or after exposure of increasing doses of UV radiation (B), in each case without induction of RNAi (left graphs) or after induction of RNAi (right graphs). Average cell density is shown at three time points and error bars denote standard deviation from three experiments.

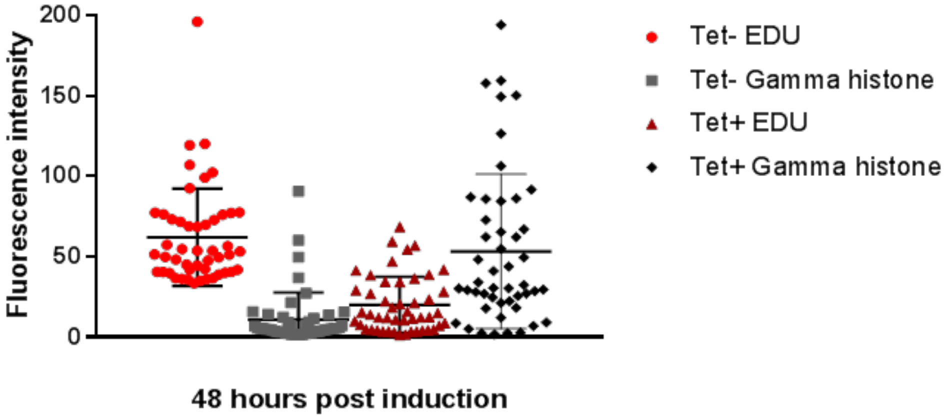


**Figure S5. EdU and γH2A fluorescence intensity after TbPolN RNAi.** Fluorescence intensity of EdU and γH2A (gamma histone) signal was calculated in Image J as the corrected total cell fluorescence (CTCF), using a region of interest (ROI, 21x21 pixels) around the cell and subtracting background signal. Dots represent the signals obtained for each individual cell, which were either induced for RNAi (Tet+) for 48 hrs, or grown for the same length of time without induction (Tet-). A total of 50 cells were analysed for each group.

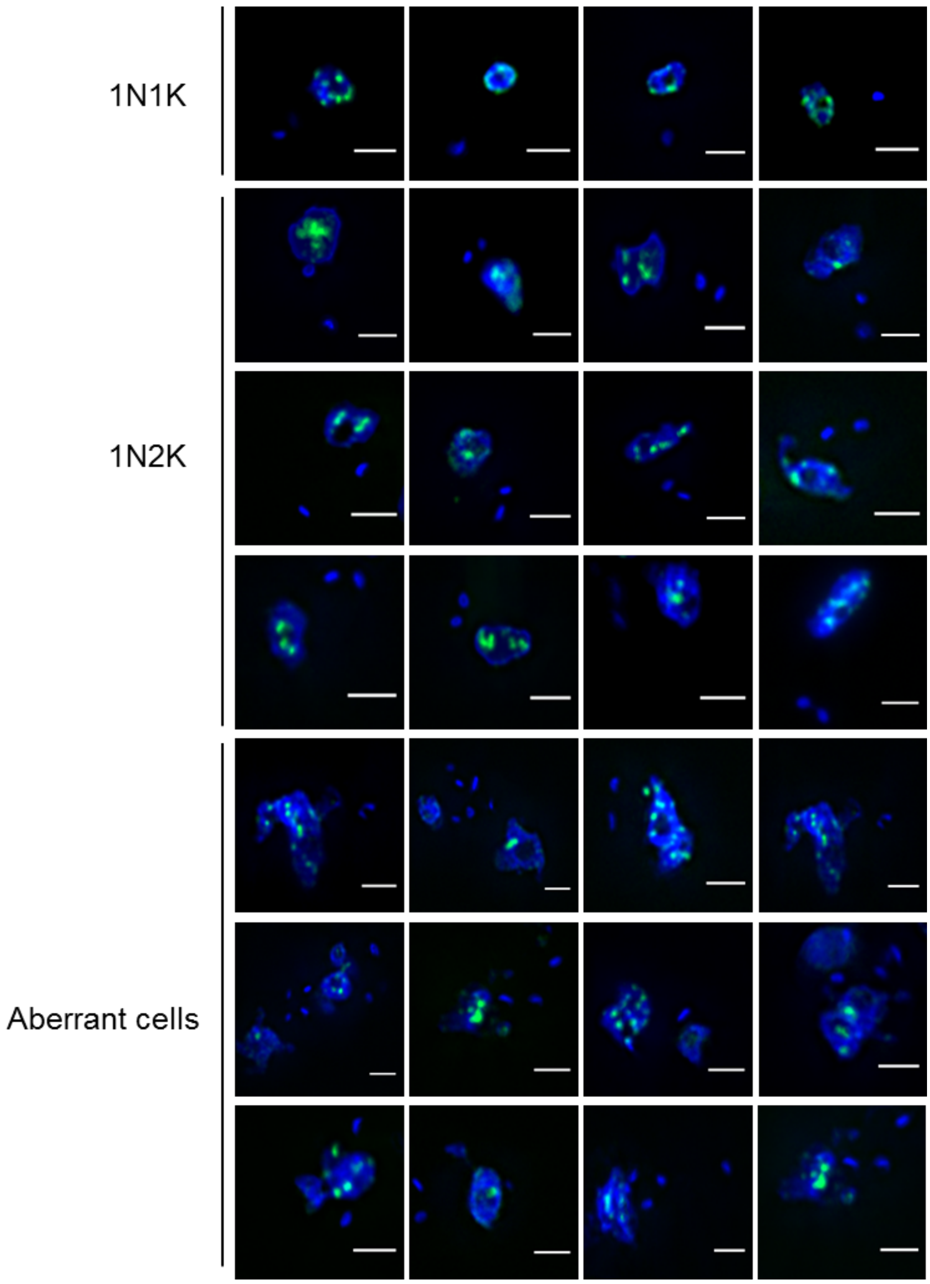


**Figure S6. Fluorescence *in situ* hybridization of telomeres 48 hrs after depletion of TbPolN.** Representative images of cells after fluorescence *in situ* hybridization using telomeres as a probe (green) and after 48 hrs growth with induction of RNAi against TbPolN. Top panels show telomere distribution in 1N1K cells, middle panels show telomere distribution in 1N2K cells, and bottom panels show telomere localisation in aberrant cells. DAPI is in blue; scale bar, 2 µm.

Figure S7

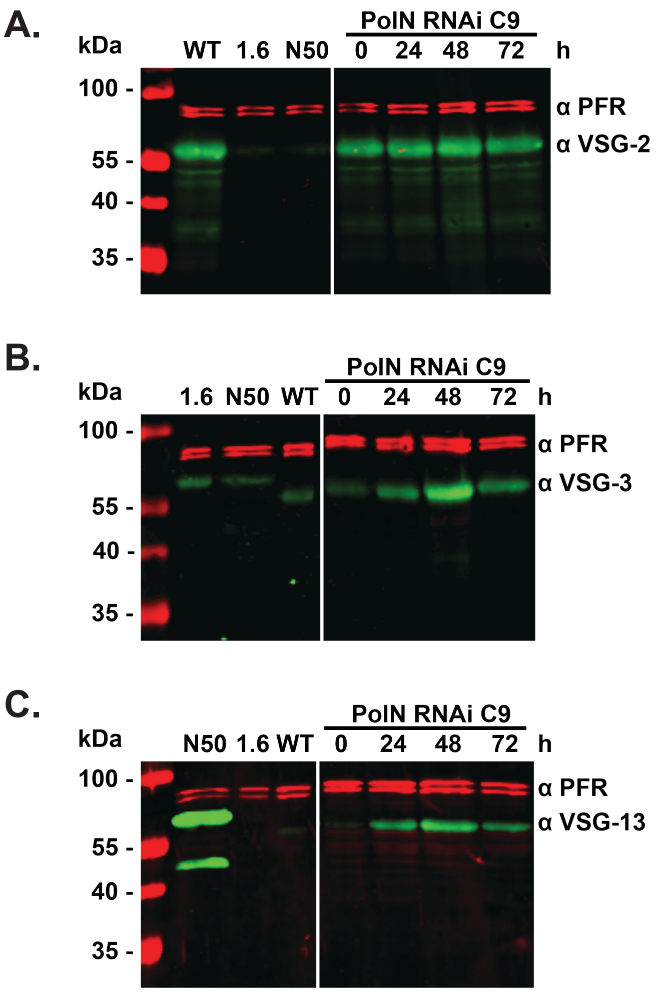


**Figure S7. Increased expression of silent VSGs after RNAi depletion of TbPolN.** Whole cell lysates were prepared at different time points after induction of RNAi against PolN in clone C9 and probed, after blotting, with antibodies specific for VSG-2 (A), VSG-3 (B) and VSG-13 (C). Antiserum against paraflagellar rod protein (PFR, red) was used as loading control. Control cell lines (left panel) were taken from Figure 10.
